# Distinct neural codes in primate Hippocampus and Lateral Prefrontal Cortex during associative learning in virtual environments

**DOI:** 10.1101/2021.08.20.457136

**Authors:** B. W. Corrigan, R. A. Gulli, G. Doucet, M. Roussy, R. Luna, A.J. Sachs, J. C. Martinez-Trujillo

## Abstract

The hippocampus (HPC) and the lateral prefrontal cortex (LPFC) are two cortical areas of the primate brain deemed essential to cognition. Here we hypothesize that the codes mediating neuronal communication in HPC and LPFC microcircuits have distinctively evolved to serve plasticity and memory function at different spatiotemporal scales. We used a virtual reality task in which animals navigated through a maze using a joystick and selected one of two targets in the arms of the maze according to a learned context-color rule. We found that neurons and neuronal populations in both regions encode similar information about the task. Moreover, we demonstrate that many HPC neurons concentrate spikes into bursts, whereas most layer II/III LPFC neurons sparsely distribute spikes over time. As the animals learned the task HPC neurons, but not LPFC neurons, increased their burst rate as a function of performance. When integrating spike rates over short intervals, HPC neuronal ensembles reached maximum decoded information with fewer neurons than LPFC ensembles. Our results show that during associative learning HPC principal cells concentrate spikes in bursts enabling temporal summation and fast synaptic plasticity in small populations of neurons and ultimately facilitating rapid encoding of associative memories. On the other hand, layers II/III LPFC pyramidal cells fire spikes more sparsely distributed in time and over a larger number of neurons. The latter would facilitate broadcasting of signals loaded in short term memory across neuronal populations without necessarily triggering fast synaptic plasticity.

## Introduction

The primate lateral prefrontal cortex (LPFC) and hippocampus (HPC) are two regions that integrate high level sensory information and play an important role in memory function. Lesion studies have shown that the hippocampus plays a fundamental role in long-term memory formation, for which hippocampal microcircuits possess enhanced synaptic plasticity (Bittner, Milstein, Grienberger, Romani, & Magee, 2017) via long-term potentiation (LTP) and long-term depression (LTD) mechanisms (Raus Balind et al., 2019). On the other hand, the prefrontal cortex, particularly its lateral aspect (LPFC areas 9/46), is known to play a fundamental role in short-term memory encoding, specifically layers II/III (Arnsten, 2013; Spaak, Watanabe, Funahashi, & Stokes, 2017). Short-term memories (e.g., shortly remembering a telephone number) are encoded in patterns of neural activity across populations of neurons that vanish after a few seconds (Leavitt, Pieper, Sachs, & Martinez-Trujillo, 2017). Therefore, it is reasonable to assume short-term memories do not necessarily trigger the mechanisms of long-term synaptic plasticity, at least within short time scales. Here we hypothesize that the neural codes underlying neuronal communication and information processing in primate HPC and layer II/III LPFC microcircuits are spatiotemporally distinct and tailored to service the corresponding memory functions of the two structures.

In the nervous system, bursts of action potentials encode and transmit information between neurons, and they can trigger changes in connectivity by inducing synaptic plasticity. A basic mechanism that links trains of action potentials to synaptic plasticity is temporal summation: a compression of synaptic events over time that produces coincidence of postsynaptic potentials and can trigger plastic changes in individual synapses (Kandel, Schwartz, Jessell, Siegelbaum, & Hudspeth, 2012). Neuronal firing patterns that cluster spikes over short time intervals such as bursts can produce temporal summation of postsynaptic potentials in target neurons and therefore induce synaptic plasticity (Remy & Spruston, 2007; Thomas, Watabe, Moody, Makhinson, & O’Dell, 1998). Burst firing in individual neurons can often be found in brain areas associated with long-term memory formation, such as the HPC (Bliss & Collingridge, 1993; Lisman, 1997). However, such firing patterns have also been reported in areas of the primate neocortex such as the prefrontal cortex classically associated with encoding short-term memories (Womelsdorf, Ardid, Everling, & Valiante, 2014). An issue that has not been thoroughly studied in primates is how the ability to fire bursts of action potentials compares in HPC and LPFC neurons during behavior.

In the primate lateral prefrontal cortex (areas 8a and 9/46) spike bursts in layer V neurons are associated with the onset of selective attention and are synchronized with the phase of beta and gamma frequencies in the anterior cingulate cortex (ACC) (Womelsdorf et al., 2014). However, during tasks involving short-term memory (also referred to as working memory in the non-human primate literature), most studies in primate LPFC have computed spike rates over a second or more, and documented the existence of persistent firing in layers II/III (Leavitt, Pieper, Sachs, & Martinez-Trujillo, 2017). Using linear classifiers some studies have shown that spike rates computed over at least 400-500 millisecond intervals maximize decoded information (Leavitt et al., 2017; Roussy et al., 2021). This may suggest that neurons in LPFC use a sparser code compared to HPC neurons.

One possibility is that in the HPC and the LPFC the time structure of the spike train is ‘tailored’ to perform different functions. In the HPC, spikes may be concentrated in bursts to maximize the probability of plastic changes in individual synapses via temporal summation. This would be in line with the primary role of the HPC in long-term memory formation and consolidation (Eichenbaum et al., 2016). In the LPFC, spikes may be more distributed over time, a sparse code, which may favour information encoded over a larger population of neurons. In the latter scenario, ensembles of neurons can temporarily maintain and broadcast short-term memory signals locally and to other brain areas without necessarily triggering fast plasticity in individual synapses. The latter would be compatible with the role of neurons in layers II/III of the LPFC in encoding short-term memory (Constantinidis & Goldman-Rakic, 2002; Fuster & Alexander, 1971).

Here, we compare the prevalence of spike train bursts in the HPC and LPFC of macaque monkeys performing a spatial navigation memory task in a virtual environment and their relationship to task performance. This type of task has the particularity of delivering a more naturalistic, and fluid nature to the behavioral trials because the animals navigate through a complex virtual environment using a joystick to perform the task and the eye movements are unconstrained. We found that neurons in both structures encode information about task variables (e.g., task periods and memory associations). However, HPC neurons more often compress spikes into bursts relative to LPFC layers II/III neurons. In the HPC but not in the LPFC bursts increase in frequency as animals learn the task and behavioral performance improves. Using linear decoders, we demonstrate that it is possible to decode task related information from burst rates in the HPC with an accuracy similar to that of decoders using spike rates integrated over similar time windows. On the other hand, in the LPFC burst rate decoders performed substantially worse than spike rate decoders. Additionally, we demonstrate that small ensembles of HPC and LPFC neurons are sufficient to provide task related information, but HPC neurons do so over shorter time windows and with fewer neurons relative to the LPFC. We analysed data from LPFC neurons recorded during a naturalistic working memory task and found that burst rates were similar to those in the associative memory task, and that rate decoding performed close to chance relative to spike rate decoding. The latter suggests that LPFC layers II/III neurons use a sparse code independently of the task.

## Results

We trained 4 monkeys (*Macaca mulatta*) on a context-object association task in a dynamic virtual environment. We recorded the responses of neurons from area CA3 of the Hippocampus (HPC) of two animals and from layers II/III of area 9/46 of the lateral prefrontal cortex (LPFC) in the other two animals. During the task animals navigated through a virtual X maze using a joystick (Fig. 1a,b). Upon arriving at a decision point, where the maze branched out into two arms, two objects appeared at the arms’ ends. The animal had to select one of the two objects to obtain a reward. The objects were two colored disks and the target object was defined by the contingency of two features, the color of the object and the texture of the walls (e.g., for objects blue and green: when the texture of the maze walls was wood the target was blue, whereas for a steel wall texture the target was green) (see Fig. 1d, methods, and (Gulli et al., 2020)). Monkeys learned a new association (target color – wall texture) every day until they became proficient at the task (see example trajectories in Fig. 1c). Different color-wall texture associations were achieved by changing the target colors while leaving the wall cues (contexts) the same across sessions. In a 50-trial performance assessment window (see methods), they achieved average performances of: 75.8 % (monkey W, HPC), 61.3 % (monkey R, HPC), 74.5 % (monkey T, LPFC), and 84.3% (monkey B, LPFC) correct trials. The theoretical chance performance in this two-alternative forced-choice task was 50%.

**Figure 1.**
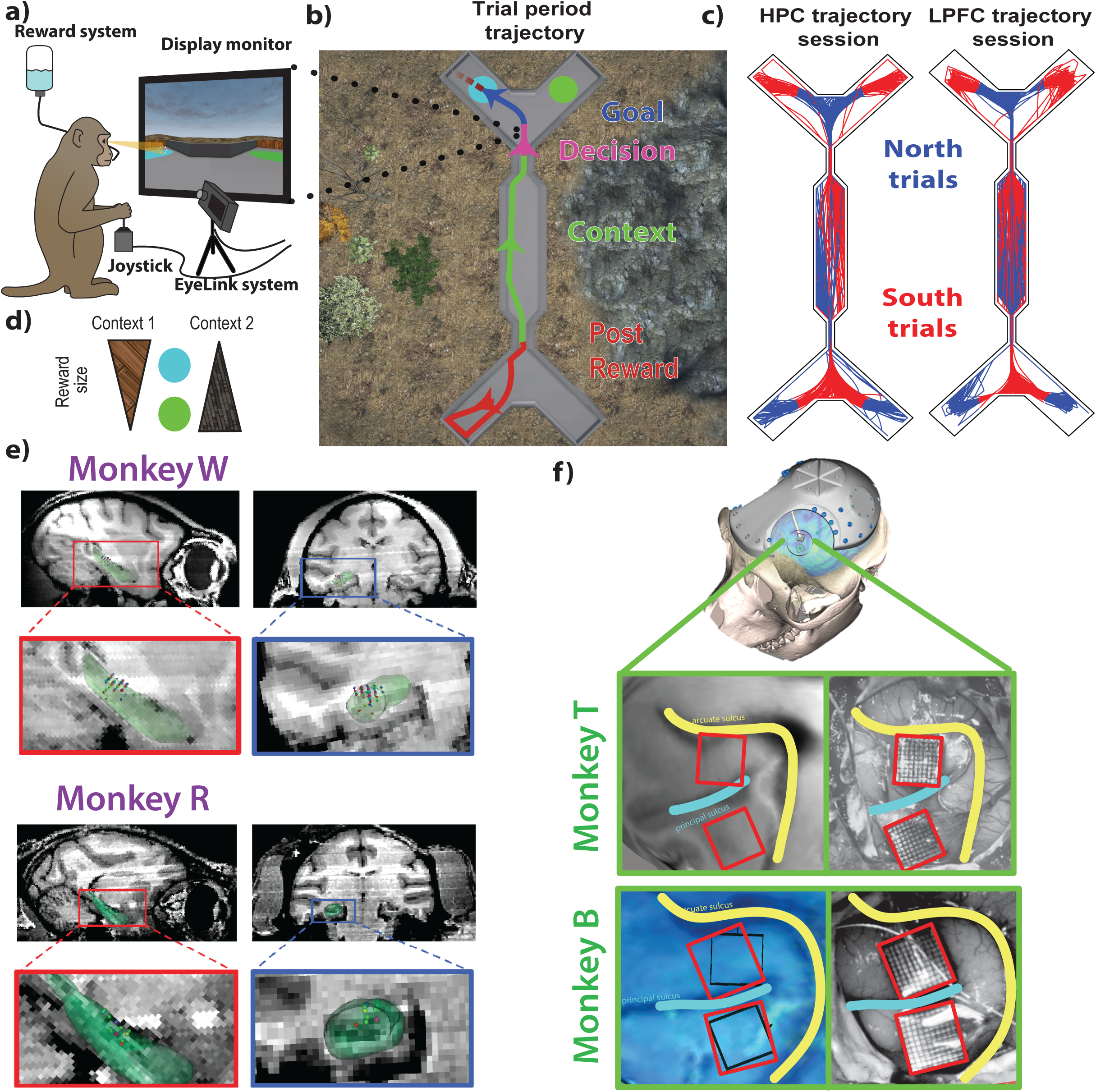
Recording from NHPs during virtual navigation learning task. a) Experimental set-up where monkeys were seated in front of a monitor and used a joystick to navigate the virtual environment. Eye position was monitored, and juice used as reward. b) Top-down view of maze with example trajectory split into four different task periods: Post reward, Context, Decision, Goal (approach). c) Example sessions from monkeys and from each area showing all trajectories separated into north and south trials. d) The rule example defined cyan as the higher value object in context one (wood), and lower in context two (steel), and the inverse for the green object. e) MRI-based reconstruction of recording positions in right hippocampus of NHPs W and R. f) Array locations on area 8a and 9/46 for NHPs B and T.

### Prevalence of burst firing in HPC and LPFC single neurons

We recorded the responses of neurons in the HPC using single electrodes (Fig. 1e) and in the LPFC using microelectrode arrays (Utah arrays, 10×10 electrodes 1.5mm electrode length) implanted in area 9/46, ventral and dorsal to the principal sulcus targeting layers II/III (Fig. 1f). Data were spike sorted and single action potential times were extracted and synchronized to the different task events. We separated neurons in each area into narrow and broad waveforms based on the difference of the trough to peak time (Torres-Gomez et al., 2020). We fit a sum of two Gaussian functions to the trough to peak distribution of recorded neurons in each area, and set the separation threshold between narrow and broad spiking at the local minimum between the two peaks (Sfig 1a,b). This method shows some inaccuracies at separating putative interneurons from pyramidal cells, for while most of the broad spiking neurons are pyramidal cells, some interneurons fire broad spikes (Torres-Gomez et al., 2020). However, considering that the majority of the neurons in the cortex are pyramidal cells (DeFelipe, 2012; Defelipe et al., 2013; Yuste et al., 2020), we restricted our analyses to broad spiking neurons and assumed that they were in their majority pyramidal cells. Pyramidal cells broadcast information between different brain regions, and in structures such as the HPC and the LPFC, they play a fundamental role in memory coding. We recorded from 205 HPC neurons over 37 sessions, and 333 LPFC neurons over 2 sessions, all of these being broad spiking. We quantified the firing rates of these neurons in both areas during the different task periods. The spikes of HPC neurons frequently occurred within 7 millisecond intervals (example on Fig. 2a, and blue spikes in c). This was not the case for LPFC neurons that fired spikes more sparsely (example in Fig. 2b, d). We additionally computed the interspike interval (ISI) distribution of the two example units per trial period, and for all periods the HPC unit shows a bias to short ISIs (Fig. 2e) compared to the LPFC unit (Fig. 2f).

**Figure 2.**
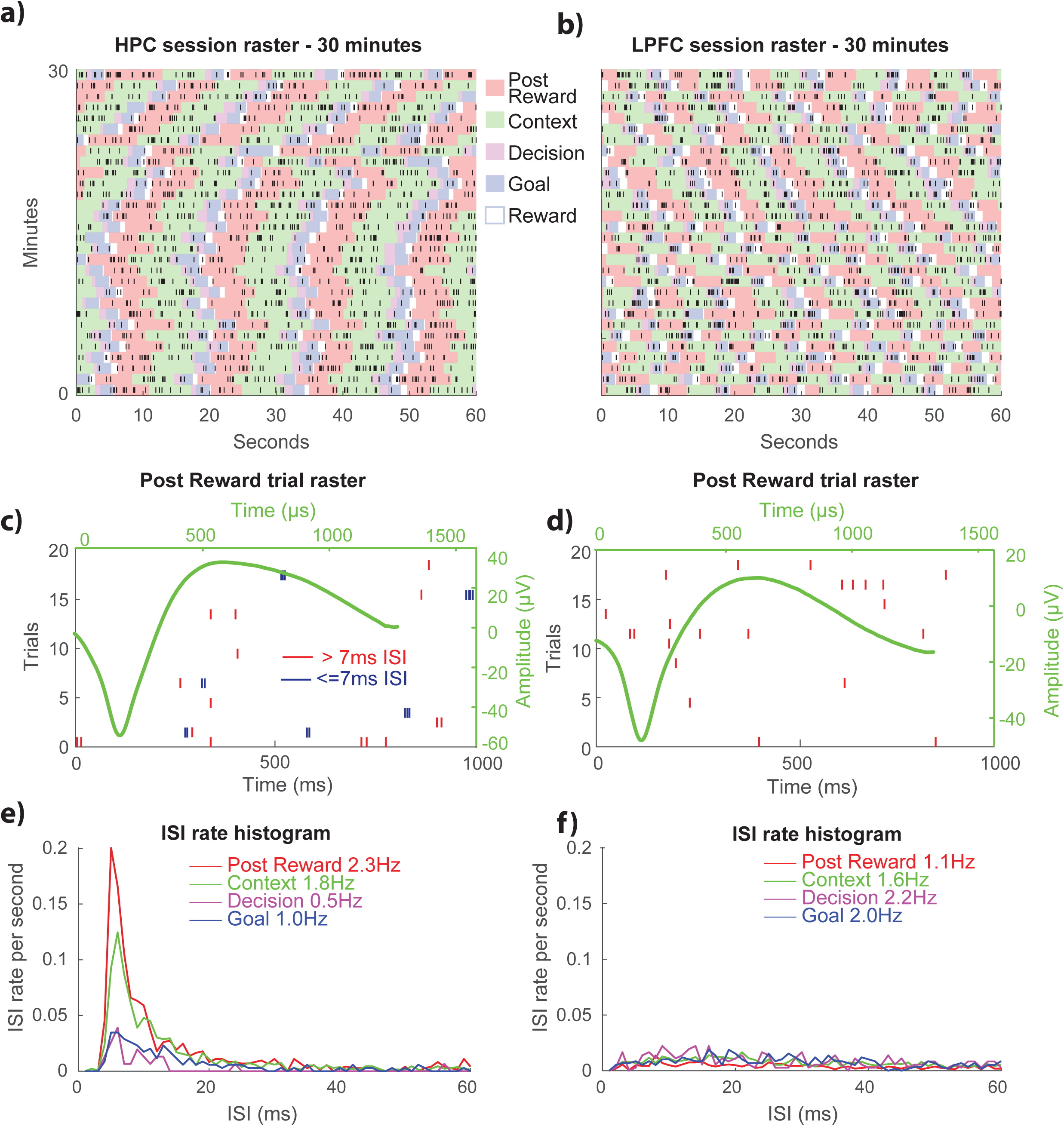
Differences in firing patterns between two example units in HPC and LPFC. a,b) Session rasters showing example unit activity for 30 minutes, with background indicating the different trial periods. c,d) Trial rasters for the post reward period with ISIs <= 7 indicated, occurring more frequently in the hippocampus. Super-imposed are the example unit’s waveform - broad spiking neurons with similar task firing rate of 1.8 Hz. e,f) Inter-spike interval (ISI) rate distributions for the different periods. Note that the hippocampal unit has much more activity below 20ms than in the following 20ms, whereas the LPFC unit has a more uniform distribution.

We examined the probability density function for ISIs across all broad spiking neurons during the period between the stat and end of the task, including the inter-trail intervals, in the two regions (Fig. 3a,b). As in the example units, HPC neurons had a large peak below a bursting threshold of 7ms ISIs compared to the LPFC. Additionally, we calculated the rate of change for ISI values in the HPC distribution and found it approaches 0 at around 7ms (fig. 3a inset). This suggests that HPC neurons frequently fire bursts with ISIs around 7ms. To further examine the differences in burst firing between the two areas for all broad spiking neurons, we measured a burst fraction which quantifies the proportion of ISIs at or below 7ms (Fig. 3c,d). HPC neurons had a significantly higher burst fraction (median = .092) than LPFC neurons (median = .027) (rank sum test, Z(216,367) = 10.45, *p<* .05) (Fig. 3c). We noticed that as firing rate increases, the probability of ISIs being below 7ms increases in the LPFC (Pearson correlation coefficient *r* = 0.67, *p* < .05). This was not the case in the HPC where the correlation was not statistically significant (*r* = −0.06, *p* > .05) (Fig. 3b). The positive correlation between burst fraction and firing rate in LPFC neurons may suggest their ISI distribution is approximated by a Poisson distribution (with some deviance as it approaches 0). The ISI distributions peak close to 0 and then decrease, and this peak rises with the firing rate because the more spikes the neuron fires in a given time window, the more frequent short ISIs become.

**Figure 3.**
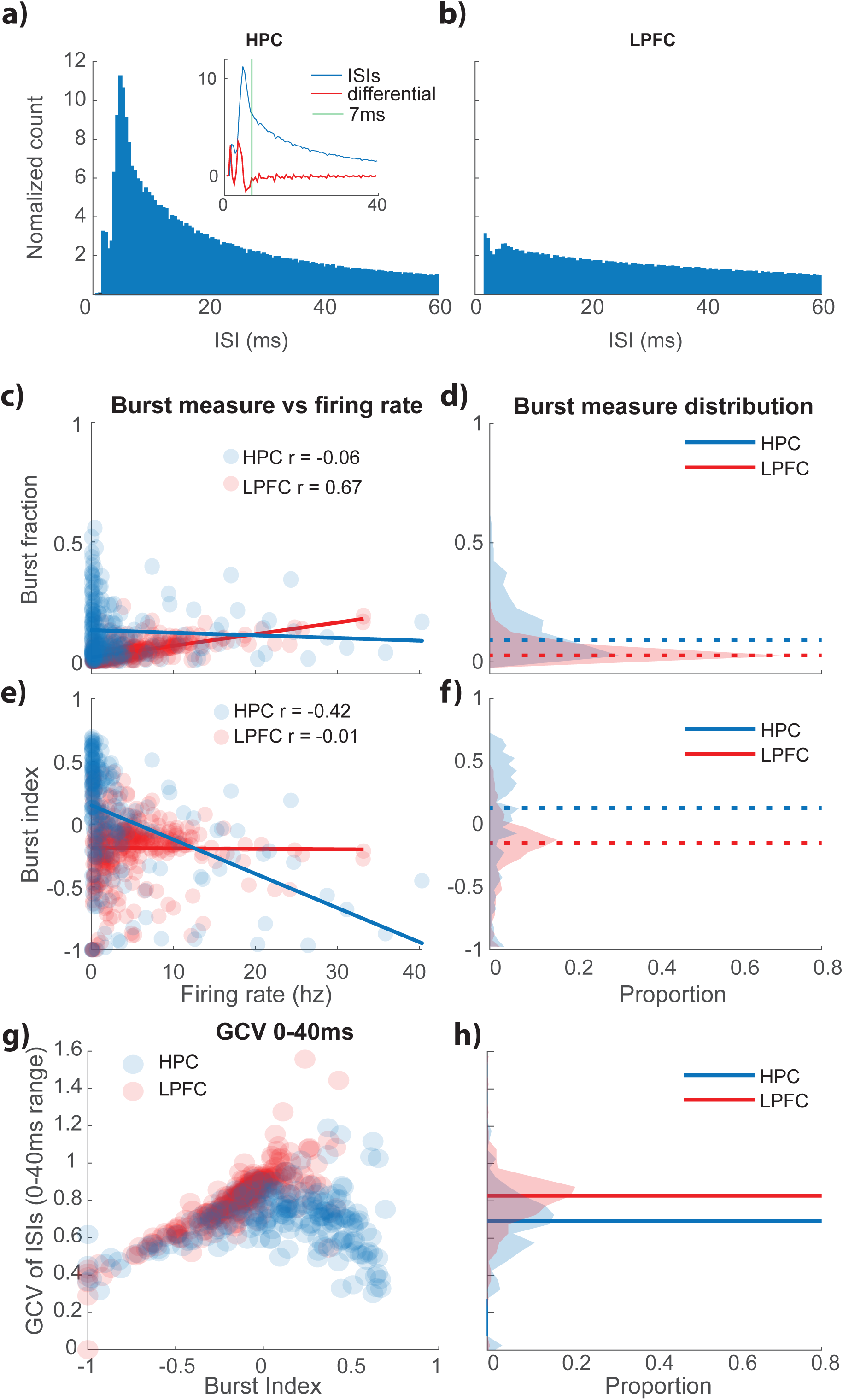
HPC cells are more prone to bursting, and LPFC burst fraction is correlated with firing rate. a) The probability density distribution of all ISIs for broad spiking neurons in the HPC, normalized to the value at 60ms. The HPC ISIs have a pronounced peak below the 7ms burst threshold. The inset demonstrates the relationship between the distribution and its first derivative, which stabilizes after 7ms. b) Same as a, except for broad LPFC neurons. c) Burst fraction plotted against firing rate, where HPC cells have high burst fraction at low firing rates. d) Population density of burst fractions. e) Same as c, but with burst index, where LPFC burst index is no longer correlated with firing rate, and there is a slight negative correlation for the HPC, see text for equation of burst index. f) Population density of burst indices. g) GCV for ISIs between 0 and 40ms plotted against burst index. HPC GCV values decrease at high burst index values.

We additionally used a method from Livingstone et al. (Livingstone, Freeman, & Hubel, 1996) to compute a probability distribution for ISIs as a function of firing rate relative to that predicted by Poisson firing neurons. We compared this predicted distribution of ISIs for a neuron using the measured firing rate over the period of the task with the actual distribution to obtain a burst index (BI, see methods). This index is a bounded ratio of the measured ISIs and the Poisson predicted ISIs below a threshold of 7ms. A positive value means shorter ISIs than predicted by a Poisson distribution, and suggests the cell often fires bursts, while a value at or below 0 indicates that the cell does not burst more than predicted for a Poisson firing neuron. We found that the BI of HPC cells (median = 0.14) were significantly higher than the BI of LPFC cells (median = −0.11, Z(216,367) = 8.60, *p<*.05) (Fig. 3e). The BI for LPFC cells did not correlate with firing rate (*r* = −0.007, *p>*.05). The BI for HPC cells was significantly negatively correlated with the firing rate (*r* = −0.42, *p<*.05) indicating that as firing rate increases BI decreases. This result seems counterintuitive. One likely explanation is that at low firing rates many spikes in HPC neurons are concentrated within bursts while at high firing rates neurons fire spikes ‘outside’ bursts. The latter leads to the same number of bursts with increases in number of spikes and consequently in firing rate.

To analyse the effect of using a certain ISI threshold to define a burst on BI, we additionally calculated the BI using different burst thresholds: 4, 7, 10, 15 and 20ms (Sfig. 2). We observed that at the 4ms threshold, very few neurons show BI values >0 in either area. For thresholds of 7ms or higher more neurons had values closer to or >0. The distributions’ median values were within a narrow span, HPC range = 0.16 to 0.25 and LPFC range = −0.1 to 0.01. We carried out the rest of the analyses using the burst threshold of 7ms because of the similar ranges found for other thresholds, and because of the stabilization of the differential values at 7ms (inset in fig 3a). Some analyses may include other thresholds when required.

To rule out the possibility that the increased bursting found in the HPC was due to spike sorting issues that may have resulted in classifying multiple units as a single unit, we calculated a signal to noise ratio measure (Sfig. 3). In brief, we calculated the difference between the trough (or peak) and the baseline voltages, and then divided this by the square root of the sum of the variance of the baseline and the trough (or peak) to get a d’ measure for both the trough and the peak (eq. 3) (Leavitt, Pieper, Sachs, Joober, & Martinez-Trujillo, 2013). We did not find a significant correlation between BI and d’ for HPC (*r* = −0.01, *p>*.05), and a small negative correlation for LPFC (*r* = −.037, *p<*.05). This indicates that there could be some influence of spike sorting noise increasing the bursts in the LPFC, but less in the HPC. These results suggest that any over estimation of burst occurrence may have happened in the LPFC data, yet bursts were more often found in HPC neurons, which argues against this variable acting as a confound to our main results.

Finally, we calculated the geometric coefficient of variability (GCV, see methods, eq. 4) for ISIs between 0 and 40ms. The GCV quantifies the variability of the interspike intervals (ISIs). If spikes are generated within bursts with regular ISIs, like in the HPC example unit (Fig. 2e), then the GCV will be low. If spikes are more sparsely distributed with variable ISIs as in the LPFC example unit (Fig. 2f) then the GCV will be high. We found that in both areas GCV increases with BI for BI values lower than zero (Fig. 3g). However, HPC neurons with positive BI values deviate from that trend (blue circles in fig. 3g): high BI HPC neurons show the lowest GCV values, as if all ISIs were clustered within a narrow range strongly indicating that bursts show highly regular ISIs and they may be at least in part influenced by intrinsic properties of these HPC neurons. On the other hand, for LPFC neurons, GCV increases as BI increases for both negative and positive BI values (Fig. 3g red circles).

### HPC bursting increases with task performance in post-reward period

We hypothesized that if bursting plays a role in learning and encoding of long-term memory, we should find a correlation between burst rates and task performance (e.g., higher burst rates should be positively correlated with performance rates). During the task, animals learned the association between the context and object colour such that their hit rate improved over the course of the session (fig 4a). To corroborate this observation, we binned the data in epochs of 20 trials evenly distributed through the session (beginning, ¼, ½, ¾, and end) and calculated the mean performance rate for each epoch and for each session. There is indeed a significant positive correlation (positive slope between performance rate and time epoch) for the 37 sessions for the HPC monkeys, and for the 17 sessions for the LPFC monkeys. The slope for the HPC sessions was 0.06 (95% CI = 0.04-0.08, Fig. 4a) and the slope for the LPFC sessions was also 0.06 (95% CI = 0.04-0.08) indicating animals learned the task at similar rates across all sessions.

**Figure 4.**
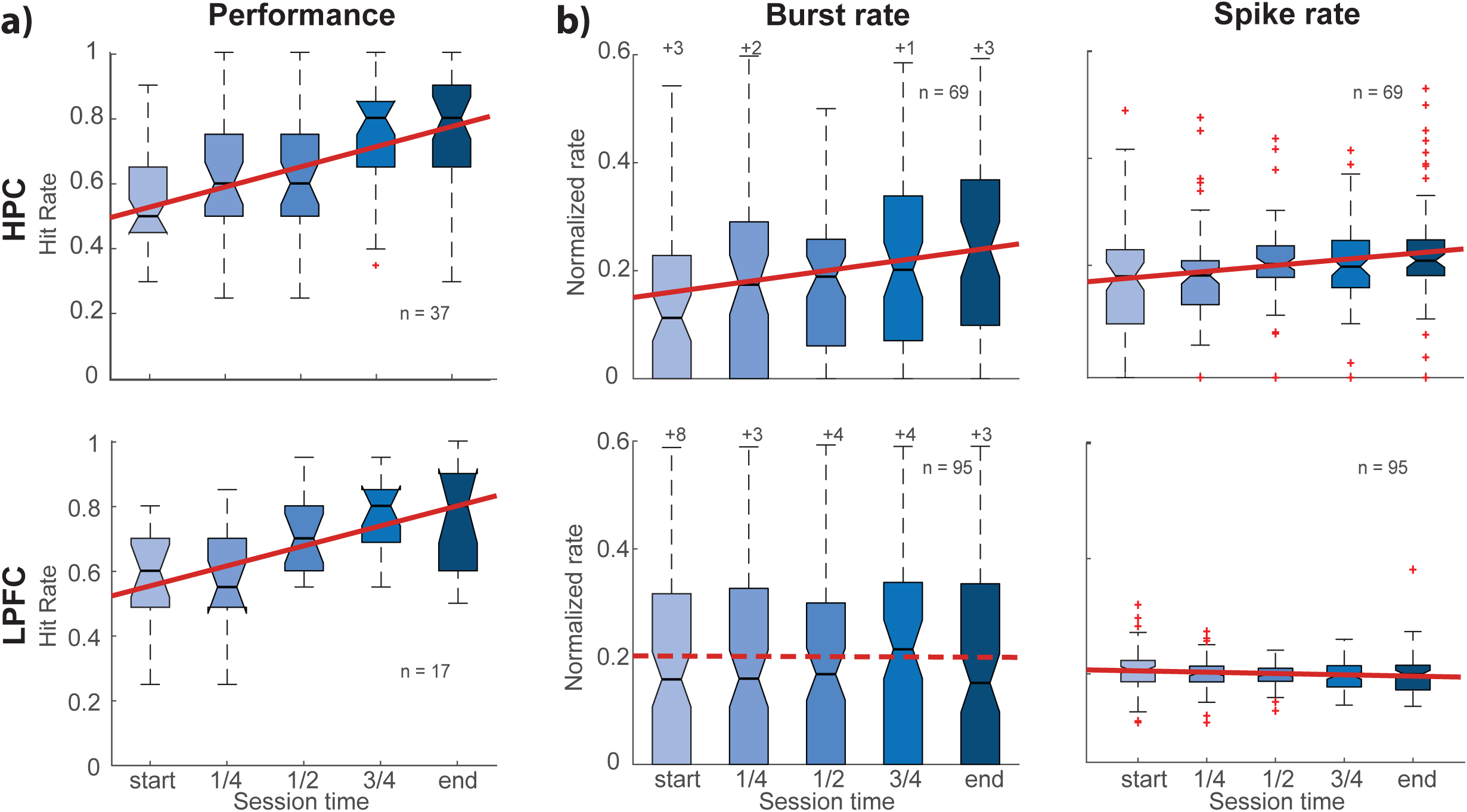
HPC bursting increases with performance, while LPFC bursting does not. a) Performance defined by hit rate for 20 trial blocks evenly spaced throughout each session for both HPC and LPFC monkeys. b) Normalized burst and spike rates for blocks of 10 correct trials during the post reward period. HPC Bursts have a significant positive slope, as do HPC spikes, while LPFC bursts are stable and LPFC spikes have a slight but significant negative slope.

To assess rates of only correct trials during the same periods, we measured the burst rate and spike rate during five epochs of 10 correct trials evenly spaced throughout each session (Fig 4b). For HPC neurons that had bursts in at least two epochs, bursting increased with a slope of 0.020 (95% CI = 0.007-0.033). The number of spikes also increased with a slope of 0.012 (95% CI = 0.005-0.020). For the LPFC, bursting did not significantly increase, with a non-significant slope of −0.001 (95% CI = −0.013-0.012), however the firing rate slightly decreased with a slope of −0.002 (95% CI -0.004- −0.001). The latter effect may be related to previous reports of decreases in spike rates in LPFC neurons when stimuli lose novelty (Asaad, Rainer, & Miller, 1998; Miller & Desimone, 1994; Wilson & Rolls, 1993). This result demonstrates that in the HPC but not in the LPFC bursting was positively correlated with improvements in performance as the animal learned the task. Importantly, these results provide evidence in favor of the role of bursting in long-term memory coding in primate HPC circuits.

One may argue that variables related to the time course of the session could confound the correlation analyses (e.g., more bursts at the beginning than at the end of the session). To address this issue, we specifically examined the relationship between bursting and performance for all trial epochs and their corresponding burst rates independently of epoch trial time. We correlated the hit rate for the 20 trial epochs with both the burst rate and the spike rate for both areas during correct trials (sample of 10 trials centred on hit rate epoch) (Sfig. 4). We found a significant correlation between HPC bursts and hit rate (*r* = 0.13, *p*<.05), indicating that the previous result was not solely due to time during the session. On the other hand, HPC spike rates were not significantly correlated with performance (*r* = 0.11, *p*>.05). In the LPFC both correlations were negative but not significant (LPFC bursts *r* = −0.4, *p*>.05, LPFC spikes *r* = −0.07, *p*>.05). This data confirms that in the HPC, but not in the LPFC, burst rates are positively correlated with performance.

### Information about task periods encoded in burst and spike rates in HPC and LPFC

Beside the associative learning component, our task had different periods corresponding to different segments of the maze (Post-reward, Context, Decision, Goal Approach and Reward). We examined whether neuronal spikes and/or bursts encoded information about the different task periods in the HPC and LPFC. For example, a neuron might respond to right turns during the Goal approach period. An alternative might be a neuron that, during the Post-reward period, integrates the reward feedback with the conjunction of the context and the chosen target in the previous trial. A deeper analyses of neuronal selectivities can be found in Gulli et al. (2020). Here we will first concentrate on overall measurements of information about task periods since our goal is to compare the information contained in burst rates (number of bursts of 3 or more spikes per second) and spike rates (number of spikes per second) (see methods). Our hypothesis is that bursts are units of information transmission, so burst rates would encode more or at least similar information about task periods as spike rates.

For each neuron we computed mutual information during the first four task periods (Post-reward, Context, Decision, Goal approach) for spike rates and for burst rates, using a permutation test for significance with 5000 permutations. The spike rate was defined as the number of spikes that occurred during the entire task period divided by the period’s duration. The burst rate was defined as the number of bursts (at least 3 spikes with ISIs < 7ms) during a task period, divided by the duration of the period. We also calculated burst rates with thresholds at 10ms, 15ms and 20ms (Sfig. 5), but most of our analyses focus on the 7ms threshold. To ensure a reasonable sample size, we only used neurons with at least 60 completed trials. Additionally, we split cells based on burst index (BI) values, with high bursting neurons (HBN) having a BI >0, and low bursting neurons (LBN) having a BI<0. The proportions of cells that were classified as HBN or LBN were different between areas. Of 192 HPC neurons, 131 (68%) were HBNs, whereas of the 333 LPFC neurons, 69 (19%) were HBNs (*z* = 10.8, *p*<.05).

From the 131 HBNs in the HPC, 71 (54%) encoded significant information about task periods; from those, 67 (94%) had significant spike rate information and 43 (61%) had significant burst rate information (Fig. 5a). From the 69 HBN in LPFC, 52 (68%) encoded significant information about task periods; from those, 50 (96%) had significant spike rate information, and 15 (29%) had significant burst rate information (Fig. 5a). The proportion of neurons with significant burst rate information was significantly higher in the HPC (61%) than in the LPFC (29%, *z* = 3.48, *p<*.*05*). From the 61 LBNs in the HPC, 70% encoded significant information about task periods; from those, 43 (100%) had spiking information, and 5 (11%) had burst rate information. These populations completely overlapped (Fig. 5b). From the 264 LBN in the LPFC, 200 (76%) encoded significant information about task periods; from those, 198 (99%) had spiking information and 25 (13%) had significant burst rate information (Fig. 5b). Unlike for HBN, the proportions of neurons with significant burst rate information for LBNs were very similar (*z* = 0.16, *p*>.05).

**Figure 5.**
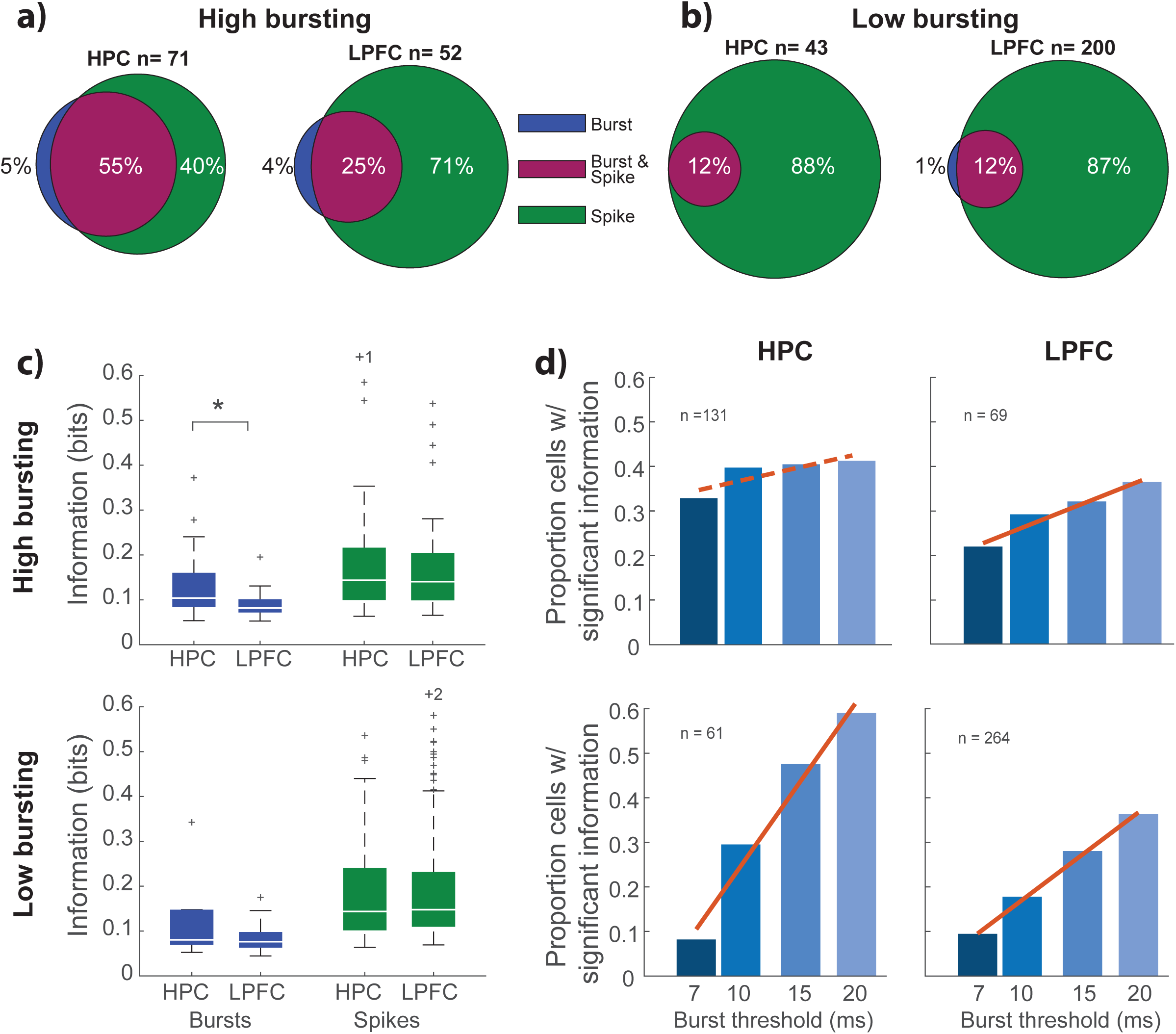
HPC high bursting neurons have more neurons with significant information, that have more bits of information than LPFC. a) Proportions of HBNs that have significant information in either the burst rate, the spike rate, or more often, both rates. b) Same as ‘a’ but for LBNs. c) Amounts of information in bits for each significant neuron in HBN (above) and LBN (below) populations. Spikes have more information than bursts in all sub-populations. HPC HBNs have more information in the burst rate than LPFC neurons. d) Proportion of cells that have significant information for different burst thresholds. Proportions significantly increase for every group except high bursting HPC.

Turning to the number of bits of information that these significant neurons had, there were not only more cells with significant information in the spike rates, but the spike rates also had more information on average in the HBNs in both the HPC (burst median = 0.10, spike median = 0.14, rank-sum, *z =* 2.48, *p*<.05) and the LPFC (burst median = 0.08, spike median - 0.14, *z* = 2.35, *p*<.05) (fig 5c). This was also the case for the LBNs in both HPC (burst median = 0.10, spike median = 0.14, *z =* 2.73, *p*<.05) and LPFC (burst median = 0.08, spike median - 0.15, *z* = 2.23, *p*<.05). When comparing information in HBNs, there was a significant difference between the HPC (median 0.10) and the LPFC (median = 0.08) using a rank-sum test (*z* = 2.04, *p<*.05). In terms of the information available in the spikes, the HPC (median = .14) was not significantly different than the LPFC (median = 0.14, *z* = 0.30, *p*>.05). For the LBNs, there was no significant difference between the information available in the two areas in the bursts (HPC median = 0.10, LPFC median = 0.08, *z* = 0.17, *p*>.05) or in the spikes (HPC median = .14, LPFC median = 0.15, *z* = 0.06, *p*>.05). Thus, for neurons that had significant information, the HPC had a higher proportion of HBNs with significant information than the LPFC, and there was more information in the burst rates of HBN in the HPC relative to the LPFC.

To further explore the effect that burst threshold might have on mutual information, we calculated mutual information for thresholds at 10ms, 15ms and 20ms. Remarkably, for the HBNs in HPC, the proportion of neurons with significant information was stable across the different thresholds, with a non-significant slope of 0.005 (95% CI = −0.007 - 0.018). In contrast, there were significant increases in the three other sub-populations: LPFC HBN (slope = .010, 95% CI = 0.001- 0.020), LPFC LBN (slope = 0.020, 95% CI = 0.014-0.027) and HPC LBN (slope = 0.038 95% CI = 0.011 – 0.065, Fig. 5d). These values are in proportions, so a slope of .01 translates to an increase of 1% of cells per millisecond added to the threshold. In general, if more neurons become significant as the threshold is increased, it is likely due to a lenient threshold, which now classifies more spikes as bursts. This results in cells with significant spike rate information, that have a more Poisson-like ISI distribution, to contain enough bursts to increase the amount of information carried in those bursts. The fact that the proportion of HBNs with significant information in HPC is relatively stable across thresholds suggests that bursting in these cells is tightly constrained, and likely influenced by the neurons’ intrinsic properties (Zeldenrust, Wadman, & Englitz, 2018). These results show that for HBNs, burst rates in the HPC carry more information about task periods than in the LPFC, indicating that HPC HBNs compress information in bursts while LPFC HBNs and LBNs distribute information more sparsely over time.

### Decoding task period information from bursts and spike rates in HPC and LPFC populations

To determine how results at the level of individual neurons generalize to populations we conducted population level analyses using linear classifiers. For the case of the HPC, recordings were done over many days, so we constructed a pseudo-population of neurons by pooling neurons and drawing the same numbers of each trial period for each neuron (Mendoza-Halliday & Martinez-Trujillo, 2017). LPFC data were recorded with microelectrode arrays and contained the original correlation structure of the population. To compare LPFC data with HPC data we shuffled the LPFC trial order, destroying simultaneity and the correlation structure of the population. Thus, the analyses use pseudo-populations of HBN in both HPC and LPFC. We trained a support vector machine (SVM) to decode the trial period based on the spike rate or the burst rate of a subsampled pseudo-population of 60 cells from each area. We selected 60 trials of each period and used the spike rates and burst rates of these periods as inputs for our decoders. We ran 5-fold cross-validation, getting one average performance, and then ran 50 subsamples to get 50 average performances. We also shuffled trial labels for each population to compute chance performances, which were at the theoretical level (∼25%).

All decoding performances were significantly different from chance, with no overlap between the confidence intervals for the actual values and the permutation values (Fig. 6a). We found that HPC burst rate decoding accuracy (median = 67.92%) was lower than spike rate decoding accuracy (median = 89.58%, permutation test, *p*<.05). We found similar results in the LPFC, burst decoding accuracy (median = 47.08%) was lower than spike rate decoding accuracy (median = 89.16%, *p*<.05). To contrast burst vs spike rate decoding between the two areas, we computed a spike-burst index: (spike rate performance - burst rate performance) / (spike rate performance + burst rate performance) (Fig. 6c). There was a significant difference between the HPC index (median = 0.12, 32% higher performance for spike rates than burst rates) and the LPFC index (median = 0.31, 90% higher performance for spike rates than for burst rates) (t-test, *t*(98) = 23.8, *p*<.05). Spike rates perform almost three times better than burst codes in the LPFC than in the HPC. Thus, although in HBN spike rates were overall more informative than burst rates in both areas, burst rates had more similar performance compared to spike rates in the HPC than in LPFC.

**Figure 6.**
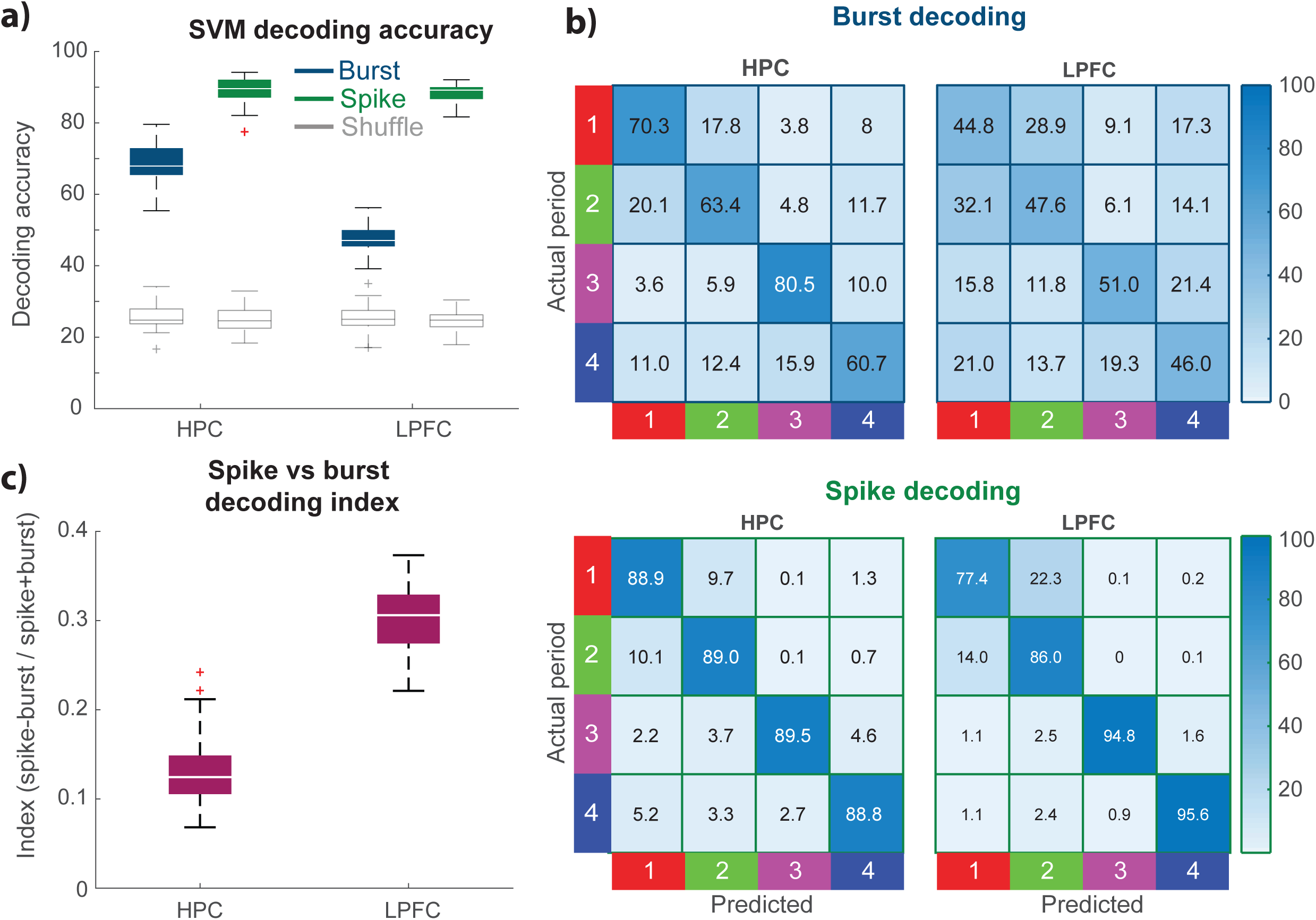
Population code for the hippocampus is similar between bursts and spikes, but not for LPFC. a) Average decoding accuracy for trial period in spikes and bursts codes in the two areas. All values are significantly different from the shuffles. b) Confusion matrices for burst and spike decoding in both areas. There is no distinct diagonal in LPFC burst decoding, but there is in the other three matrices. c) Spike-burst decoding index, where the difference between spike and burst decoding is significantly higher for LPFC, and close to 0 for HPC.

Although unlikely, it could be the case that the LPFC burst decoders performed worse than in the HPC because of differences in the number of periods without any activity (burst rates of 0 in LPFC). To assess this, we calculated the percentage of periods in each session during which a neuron was silent, for both bursting and spiking activity (Sfig. 6). We found that most neurons had a large percentage of periods without bursting activity, with the HPC having a median of 97% and LPFC having a median of 98%, which was not significantly different using a rank-sum test (*z* = 1.31, *p*>.05). For spiking activity, the HPC had a significantly higher proportion of silent periods with a median value of 55%, while the LPFC had a median of 7% (*z* = 6.43, *p*<.05). This demonstrates that there was not a significant difference in the proportion of ‘silent’ periods between areas for bursts. Additionally, the high decoding accuracy of the HPC spike rates becomes somewhat surprising considering how many more silent spike rate periods there were compared to the LPFC.

One question concerning ensemble coding of task periods from burst and spike rates is whether information encoded is distributed over many neurons or over few neurons only. To clarify this issue, we examined how decoded information from both burst and spike rates changes as a function of neuronal ensemble size and composition. For these analyses we used the HBNs that had significant mutual information in their burst rates. Using a procedure similar to Leavitt et al. (2017), we first estimated decoded information in individual neurons, and then we paired the best neurons with every other neuron to find the best duo, grouping that duo with every other neuron to find the best trio etc. (see (Backen, Treue, & Martinez-Trujillo, 2018; Leavitt et al., 2017)). We ran this process 50 times to generate a population we could analyse further. For both the HPC and the LPFC, ensembles of a relatively small number of neurons seem to saturate the decoder’s performance (Fig. 7a). To quantify this, we fit an exponential function to the data (eq. 6). All fits had an r^2^>.95. We calculated the point at 95% of the maximum performance to conduct comparisons between areas and burst and spike rate decoders. The HPC reached the 95% rate for spike rates at an average of 4.85 (SD = 1.30) neurons, and for the burst rate at a similar number of 4.74 (SD = 1.25) neurons (*t*(95) = 0.43, *p* > 0.05). For the LPFC, firing rate ensembles reached the 95% point at an average of 4.50 (SD = 1.63) neurons, and burst rate ensembles at a similar number of 4.20 (SD = 1.72) neurons (*t*(97) = 0.90, *p* > 0.05). The 95% point was not significantly lower for the LPFC than the HPC for firing rate decoders (*t*(96) = 1.18, *p* > 0.05), and there was no significant difference between the 95% points for the burst rates between areas (*t*(96) = 1.79, *p*>.05). Thus, ensembles with small numbers of neurons (∼5) were sufficient to saturate the decoder’s performance for both burst rates and spike rates.

**Figure 7.**
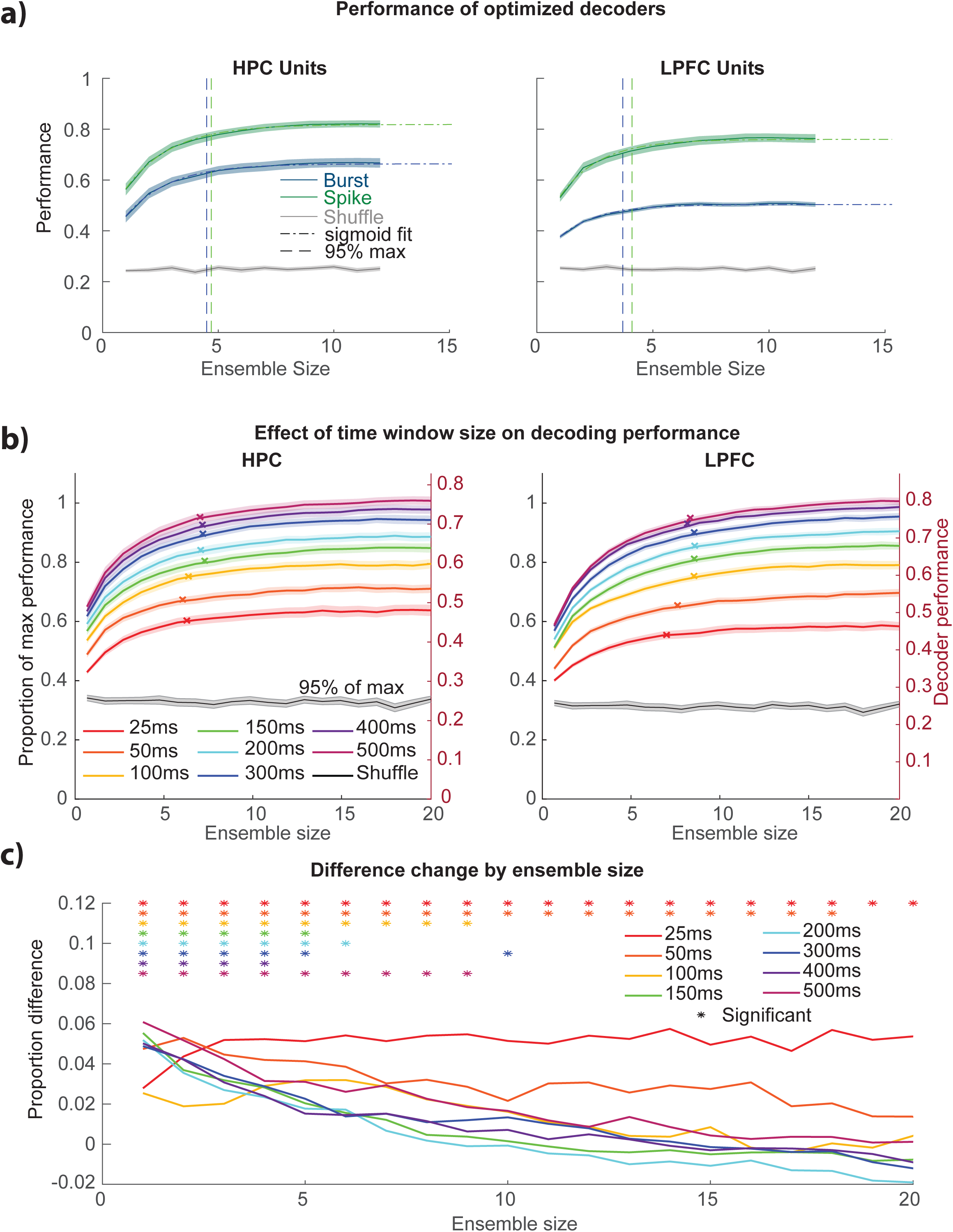
Hippocampal firing rates carry information in compressed time windows and with fewer neurons. a) Optimized ensembles for burst and spike rates yielded by neurons that have significant burst information and their fits to y = 1-a*e^(-(x/b)) -c. HPC burst rate ensembles are closer to the performance of HPC spike rate ensembles in terms of maximum performance and in terms of position of the 95% threshold crossing. b) Optimized decoders for spike rate in different time windows, where HPC ensembles have a smaller range. Left axis is performance as a proportion of maximum achieved performance and right axis is the raw performance. 95% of maximum point is indicated with an ‘x’ on each curve. c) HPC proportion of maximum performance minus LPFC proportion of maximum performance. The difference shows that for HPC, smaller ensembles perform better, as well as performing better in smaller time windows. Asterisks indicate significant difference from 0 (p <.05).

To quantify the differences in performance between burst rate decoders and spike rate decoders in each area we computed a performance index. We used asymptotes of the performance equation for all of the 50 optimized decoders. For each decoder we subtracted the burst rate performance from the spike rate performance and divided by the sum of the performances (eq. 5). The HPC performance index was lower (mean = .16, SD = .09) than the LPFC index (mean = .24, SD = .11; *t*(98) = 4.40, *p*<.05). These results indicate that burst rate contributes significantly more to information decoded from neuronal populations in the HPC than in the LPFC. Importantly, in both structures, available information can be decoded from small ensembles of neurons suggesting that small subpopulations are sufficient to encode period information during our virtual navigation task.

### Effect of integration time windows in decoded information

The previous results show that the HPC concentrate spikes in burst more often that LPFC, however, in both structures rates codes were more informative than burst codes suggesting that regarding information encoded spikes inside and outside bursts matter. To study the time scales of integration more closely for all spikes, we examined decoding performance as a function of length of the integration time window for both the HPC and the LPFC. The optimized ensemble building technique was applied to the firing rates of neurons that had significant spike rate information. However, instead of integrating over the whole task period, we took different sized time windows centered at the middle of the period (i.e., 25, 50, 100, 150, 200, 300, 400, 500ms). We repeated the same curve fitting procedure as in figure 7a and calculated the 95% point for all the average curves (Fig. 7b). The curves in each area were normalized to the maximum asymptotic performance, which happened to be at the longest window (500ms).

We found that in general, the 95% points for the HPC (mean = 7.0, SD = 0.45) were significantly lower than for the LPFC (mean = 8.3, SD = 0.56; paired t-test, *t*(7) = 8.82, *p*<.05) (Fig. 7b). This indicates that generally for the different time window lengths, the HPC seems to use fewer neurons than the LPFC to reach the asymptotic performance. We further calculated the performance of each ensemble as a fraction of the maximum of the fit of the curve for the average data in the 500ms ensembles. The latter allows subtracting LPFC fractions from the HPC fractions for every time window and ensemble size to create an ensemble index (HPC-LPFC). This inded compensates for absolute differences in maximum performance between areas. A number higher than zero means the HPC achieves a higher proportion of the maximum decoding performance with the same ensemble size. A negative number means the opposite.

To determine whether ensemble size and time window affect decoding differently in the two areas we conducted a 2-way ANOVA on the ensemble index, which was significant for both factors, time window size (*F*(7) = 65.8, *p*<.05) and ensemble size (*F*(19) = 17.6, *p*<.05); the interaction was not significant (*F*(133) = 1.0, *p*>.05). We further carried out two-sided t-tests to determine which differences between areas were significantly different from 0, indicating that an ensemble of size *n* at a particular timepoint performed better in one area than the other (*p*<.05). As demonstrated in figure 7c, at lower ensemble sizes across all time windows the HPC ensembles outperform the LPFC ensembles. Remarkably, for the 25ms time window, the HPC continues to outperform the LPFC for all ensemble sizes. These results demonstrate that HPC ensembles can ‘compress’ more information into smaller time windows (25ms) and using fewer units (up to 4) as compared to the LPFC.

### Layers II/III LPFC neurons do not increase burst rates during a working memory task

It may be possible that the differences in bursting frequency and information coding between areas is due to the fact that the associative memory-navigation task was tailored for the HPC circuits rather than for the LPFC. Thus, if the LPFC is tested with another type of task, such as a working memory task, bursting may emerge. We tested the same two animals with MEAs in the LPFC in a working memory task in a virtual environment. The monkey was placed at the start location indicated in figure 8a at the side of a virtual circular arena. During the Cue period, one of 9 possible locations (arranged in a square array) would be cued with a red fog spotlight, the cue disappeared and was followed by a 2 second Delay period where the monkey had to maintain the location in working memory. Right after, the animal was allowed to navigate to the remembered location (response period). Further analyses of performance and other details can be found in Roussy et al. (2021). Over 3 sessions (1 from monkey B, 2 from monkey T) we recorded 176 units. We assessed the bursting characteristics of the neurons recorded in the working memory task using similar analyses as in the other task.

**Figure 8.**
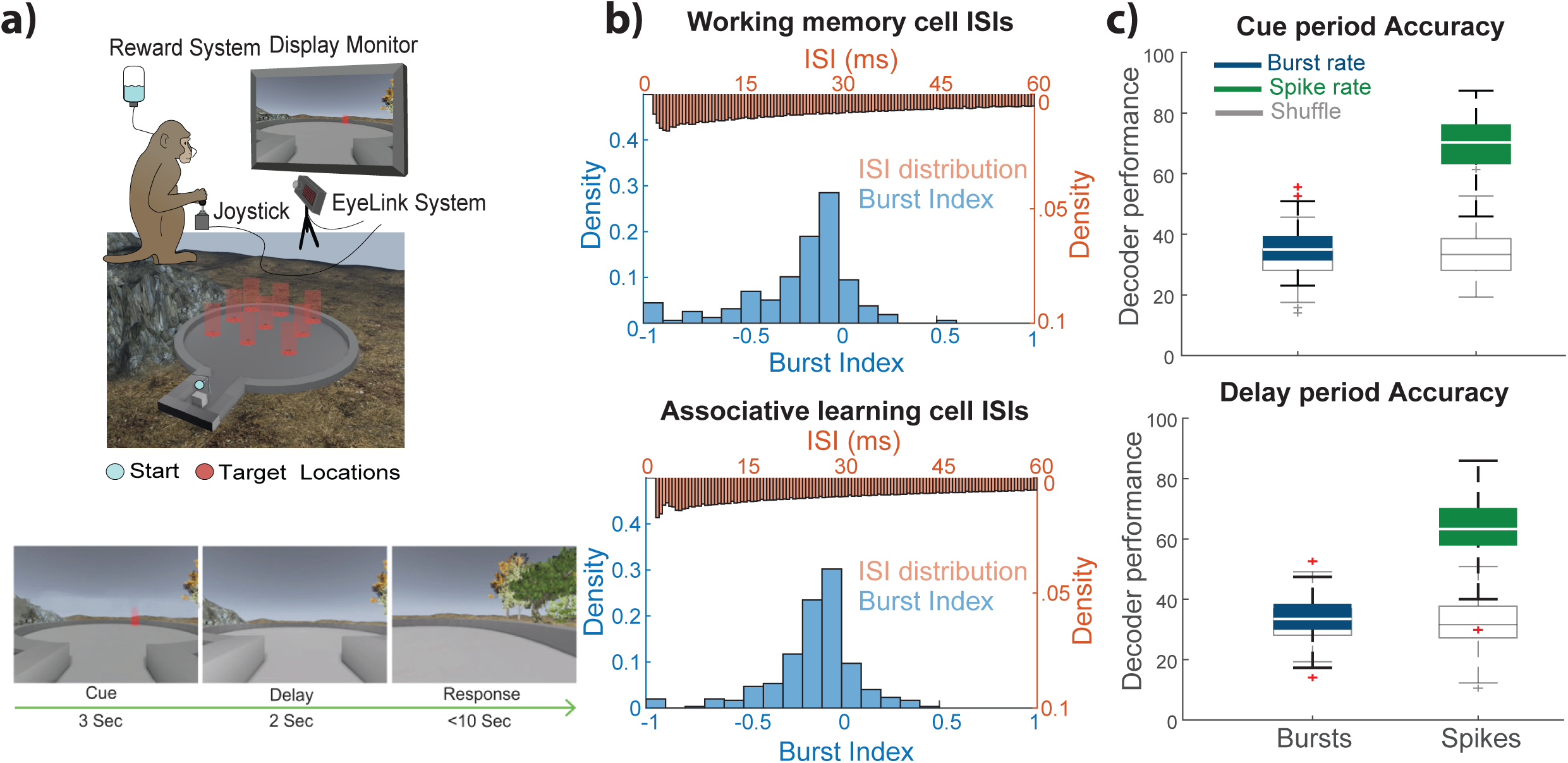
LPFC does not use a burst code for spatial working memory. a) Set-up for virtual spatial working memory task. Monkeys were again seated in front of monitor and used a joystick to navigate. Middle image has potential target locations indicated in red. Below are still images of the screen during an example trial. b) Bursting is similar for LPFC cells during the working memory task and the learning task from previous analyses, with no significant difference between burst index value distributions. c) SVM decoding performance is approximately chance for both the Cue and Delay period, while spike rate decoding is approximately double chance.

First, we found that the burst index distribution was not significantly different from the one in the associative memory task (Fig 8b, Kolmogorov-Smirnov test, *d* = 0.07, *p*>.05). We used an SVM to decode in which of the three array columns (left, middle, right) the target was positioned during both the Cue period and the Delay period. Again, we shuffled the trial labels across neurons to have an estimate of chance performance (∼33% or chance decoding) to determine significance. We analyse data from 50 different decoders of 19 neurons each. Decoding accuracy for spike rates in both the Cue (median = 70.3; permutation test, *p*<.05) and the Delay (median = 63.3%, *p*<.05) were significantly higher than chance. However burst rate decoding was not higher than chance during either period (Cue median = 35.0%; permutation test, *p*>.05, Delay median = 33.5%, *p*>.05). These results were similar to those in the associative memory task indicating that neurons in layers II/III of LPFC preferably use sparse codes over bursts to encode the contents of visuospatial working memory. Importantly these results indicate that relative to the associative memory task, bursting does not emerge in LPFC neurons when using a working memory task.

## Discussion

We tested four macaque monkeys during a virtual navigation task and recorded neuronal activity from the HPC (2 monkeys) and the LPFC (2 monkeys). We found that neurons in both areas encoded similar information about the task periods, however, HPC neurons fired bursts more often than LPFC neurons. In the HPC but not in the LPFC burst frequency was positively correlated with the animals’ performance in an associative memory task. Using linear decoders, we found that bursts were more informative about the task in the HPC than in the LPFC. We also show that HPC neuronal ensembles encode more information about that task over windows as short as 25ms and using fewer neurons relative to LPFC ensembles. Finally, we showed that bursting rates as well as the information they encode did not change in LPFC neurons when animals were tested in a virtual working memory task. In the latter scenario spike rates, but not burst rates, allowed accurate decoding of remembered locations.

### Burst firing in HPC and LPFC neurons

The time compression of spike trains in bursts has been previously studied in different model systems (Zeldenrust et al., 2018). Here we have used the definition of bursts as trains of action potentials that occur with close temporal proximity (< 7ms) or are more concentrated in time than predicted by a Poisson process. Using these criteria, we found that the HPC has a larger proportion of putative principal neurons firing bursts than layers II/III of the LPFC. This is in line with previous studies in the HPC that have reported an abundance of burst firing in HPC principal cells across species (Bliss & Collingridge, 1993; Lisman, 1997; Skaggs et al., 2007; Xu et al., 2012). It also agrees with studies in layers II/III of the LPFC reporting that information decoded during short-term memory and attention tasks is maximized when using time windows of 400ms or longer (Backen et al., 2018; Leavitt et al., 2017; Tremblay, Pieper, Sachs, & Martinez-Trujillo, 2015). We should make clear that in both structures, rate codes that count all the spikes inside and outside bursts were more informative than codes counting the number of bursts (burst codes). The latter suggests that not all informative spikes, even in the HPC, undergo temporal compression in bursts, but they exist outside the burst. Thus, the function of bursts may not be information coding per se but may be related to other functions such as triggering rapid and effective synaptic plasticity (Zeldenrust et al., 2018). Indeed, the fact that we found burst rates in the HPC, but not in the LPFC, to increase with performance during the post reward period of the associative memory task could be indicative of the role of bursting and temporal summation triggering relatively fast changes in synaptic plasticity in the HPC (Harris, Hirase, Leinekugel, Henze, & Buzsáki, 2001). Such changes may also occur in the LPFC; however, they are either triggered by other mechanisms (e.g., spatial summation) or occur over longer time periods compared to the HPC.

Our results are unlikely due to differences in the information encoded by the HPC and LPFC. We designed a task in which we could measure neuronal responses in both areas during different task periods (see Figs. 5,6). Previous studies have shown that both the HPC (Doucet, Gulli, Corrigan, Duong, & Martinez-Trujillo, 2020; Gulli et al., 2020) and the LPFC (Roussy et al., 2021) encode spatial information during navigation tasks. Additionally, both regions encode information about stimuli in an associative learning task, with information changing with task period (Brincat & Miller, 2016; Gulli et al., 2020). Indeed, our data show that single neurons and neuronal populations in both areas similarly encode task period information. This information may be related to contingencies of the task (e.g., associative learning), spatial position during virtual navigation or a combination of both (Sfig. 7). One may argue that the differences in the number of neurons firing bursts between the two areas may be due to differences in the pattern of eye movements that may have been different between the HPC and the LPFC animals. For example, one may here consider that in both areas, saccades may be preceded by a presaccadic burst and more saccades may have led to more bursts. In our naturalistic task, where animals could freely make saccades, we found that the durations of intersaccadic intervals, or foveations, is similar across HPC and LPFC sessions arguing against the explanation that saccade patterns could be the cause of the differences in burst firing (Sfig. 8). It may also be possible that cells in the different areas encode saccade parameters linked to burst firing distinctively. However, a recent study has reported that during a naturalistic task only 2-3% of LPFC neurons are tuned for saccade parameters (Roussy et al., 2021). In the HPC, saccades produce phase resetting of LFPs, but HPC neurons are in general not tuned for saccade parameters (Doucet et al., 2020; Gulli et al., 2020).

It has been reported that a feature of HPC principal cells is they frequently fire in very rapid bursts, measured by ISIs below 20ms (Lisman, 1997; Skaggs et al., 2007), or more specifically 6-8ms (Buzsáki, 2015; Ranck, 1973). This physiological burst can make up a significant proportion of all spikes, and a high burst fraction (fraction of all ISIs that are below 20ms) is a distinguishing feature of these HPC cells in primates (Skaggs et al., 2007) as well as rodents (Lisman, 1997). The hippocampal spike burst has consistently proven intriguing to scientists (Kepecs, Wang, & Lisman, 2002; Lisman, 1997; Zeldenrust et al., 2018). For example, bursts with short ISIs <6-8ms are quite common along the perforant pathway, from the dentate gyrus to the subiculum regions (Mizuseki, Royer, Diba, & Buzsáki, 2012; Pernía-Andrade & Jonas, 2014; Simonnet & Brecht, 2019). It is currently a matter of debate whether bursts can be the units of information encoded in the HPC (Harris et al., 2001). Our results in the HPC indicate that there is information carried by spikes outside the bursts. We could decode task period information from burst rates with a performance much higher than chance. However, when analyzing information decoded from bursts and spike rates (counting spikes within and outside the burst), the latter outperformed the former, suggesting that spikes outside a burst add to information coding. Importantly, in the LPFC burst decoding was much lower than spike rate decoding, suggesting that in this area spikes outside a burst play a larger role in information coding relative to the HPC.

In the macaque LPFC, few studies have examined the role of burst firing in the coding of working memory signals (Constantinidis & Goldman-Rakic, 2002; Voloh & Womelsdorf, 2018; Womelsdorf et al., 2014). Constantinidis and Goldman-Rakic used an unbounded burst index (comparing ISIs with predictions from a Poisson process) in LPFC neurons, and found the median to be .9, just below 1 which would match the Poisson prediction. They calculated their index during the fixation period of an oculomotor delayed response task, so the visual stimulation and mental state were likely quite different from the monkeys in a task like ours, with a dynamic virtual environment. This could explain why they found 10% of cells with an index value ≥4, which would be approximately .6 in our bursting index, but we found no LPFC neurons above .5. We also used a slightly larger window of ISIs, so this may have affected our results if ISI peaks drastically dropped off before 7ms. Additionally, we were restricted to layers II/III of LPFC, and their study may have included neurons in deeper layers that can be more prone to bursting. An example of bursting in deeper layers is presented in work by the Womelsdorf group in a similar cued attention task, which found that ISIs ≤ 5ms increased during selective attention in LPFC (Womelsdorf et al., 2014). They also found that narrow waveform putative interneuron bursts were synchronized to different LFP phases during attention. These results are not necessarily in contradiction with ours; we did find neurons firing bursts in the LPFC. However, the majority of the informative spikes in LPFC neurons we recorded from occurred outside the burst. Moreover, bursts were not informative when burst rates were used to decode information about task period or contents of working memory. Overall, our results indicate that bursts are less prominent and play a lesser role in information coding in layers II/III of LPFC compared to the HPC.

### Memory functions of HPC and LPFC and relationship to bursts

A main hypothesis in our study was that the features of the neural codes in the HPC and LPFC have evolved to serve the functions of these two structures. We found that bursts are more prevalent in the HPC, but also that they increase in correlation with task performance (Fig. 4) during the period where the previous trial information needs to be consolidated (and when neurons show conjunction encoding of the previous trial information (Gulli et al., 2020)). Given the relationship between burst firing, temporal summation and synaptic plasticity, our results indicate the prevalence of burst firing in the HPC serves the function of this structure in the formation of long-term memories. Indeed, bursting has been shown to be more reliable than single spikes at producing postsynaptic potentials (Remy & Spruston, 2007; Thomas et al., 1998). Xu and colleagues (Xu et al., 2012) ablated excitatory post-synaptic potentials triggered by single spikes outside bursts by knocking down synaptotagmin-1 in rodent HPC CA1. Their manipulation preserved the potentials triggered by burst spikes. Under these circumstances, learning was preserved. However, when they repeated the manipulation in the medial prefrontal cortex, learning was impaired. Although these experiments were conducted in rodents, their results suggest that in the HPC, burst spikes play a fundamental role in memory formation while spikes outside the burst play a lesser role. The inverse is true for the prefrontal cortex of the mouse.

In the LPFC, spike rates integrated over long time intervals (>400ms) provide the most information about the items held in short-term memory (Leavitt et al., 2017; Roussy et al., 2021). Indeed, when we used the 2s delay period, our decoders achieved high accuracy with spike rates, but when we used burst rates, we were unable to decode any information stored in working (short-term) memory (Fig. 8). We reasoned that this sparser code relative to the HPC may avoid triggering rapid synaptic plasticity that would lead to the consolidation of short-term memories as long-term. This would be counter-productive since most short-term memories do not need to be stored long term. It would be wasteful for the brain to trigger long-term memory storage mechanisms every time a short-term memory is ‘loaded’ in the LPFC microcircuits. Nevertheless, we did find HBNs in the LPFC layers II/III, in both the learning and the working memory task. These neurons may serve other roles not necessarily related to long-term memory formation, such as broadcasting and communication with other brain areas (Womelsdorf et al., 2014).

### Two distinct architectures for two different memory functions

Differences in the neural codes employed by the HPC and LPFC may be due to different cortical architectures that have evolved to serve different functions. Indeed, there are anatomical differences between the Hippocampus and the LPFC. The Hippocampus contains three cortical layers (paleocortex) and a relatively well-mapped circuitry in terms of input and output pathways (O’Keefe & Nadel, 1978). The HPC Cornu Ammonis (CA) subfields have limited connectivity with areas of the neocortex. The principal cells’ axons follow a well-defined connectivity pattern that ends in the subiculum. Output from the HPC is almost exclusively through the subiculum and the entorhinal cortex. On the other hand, The LPFC is extensively connected to many other brain structures (Yeterian, Pandya, Tomaiuolo, & Petrides, 2012). The LPFC is organized in 6 cortical layers with a major expansion of layers 2/3 where neurons encoding working memory have been identified (Arnsten, 2013; Constantinidis & Wang, 2004). Neurons in this region have extensive functional connectivity with one another that depends on their receptive and memory field location (Leavitt et al., 2013) and such fields cover the entire visual space (Bullock, Pieper, Sachs, & Martinez-Trujillo, 2017).

One interesting question would be whether *in vivo* burst firing regimes in the HPC are mainly due to intrinsic properties of the principal cells or to network dynamics. Studies in rodents have shown that CA3 neurons also fire bursts *in vitro* when isolated from the rest of the brain (Mizuseki et al., 2012; Ranck, 1973; Traub & Wong, 1981). This bursting behaviour may be required to induce processes such as long-term potentiation (LTP) that shapes synaptic strength and enables coding of long-term memories (Bliss & Collingridge, 1993). This suggests burst firing is robustly ‘built up’ into the intrinsic machinery of the HPC principal cell (e.g., membrane channel composition) rather than implemented into the HPC networks’ dynamics. Such intrinsic cellular machinery would make the HPC a structure intrinsically built for burst firing that enables the processes underlying long-term memory formation (i.e., changes in synaptic strength that must occur effectively and rapidly in order to engrave memories into the network structure).

In the LPFC, a recent study has reported the existence of pyramidal cells in layers II/III that are intrinsically ‘bursty’ based on their response profiles to square wave current pulses (González-Burgos et al., 2019). Interestingly, this study reported that burst neurons were more abundant in the LPFC relative to the lateral intraparietal area (LIP). Thus, it is possible that neocortical areas also differ in the way they cluster informative spikes in time depending on their involvement in long term-memory formation or their connectivity with the HPC and other areas. The latter may indicate that neural codes are heterogenous and can all serve information transmission, but at the same time they can serve different functions by changing the spatiotemporal features of spike trains. The features that make burst codes different across areas may be part of the intrinsic make up of individual cell types. The latter does not take away from the role of network connectivity and dynamics in shaping the neural code but can add heterogeneity and efficiency to neural communication and plasticity across the cortical mantle.

## Methods

Four male rhesus macaques (*Macaca mulatta*) were used in these experiments, two in HPC experiments (7 and 14 years old, and 7kg and 12kg respectively) and two in LPFC experiments (10 and 9 years old, 12kg and 10kg respectively). They were trained on a context object association task in a virtual environment where they received juice rewards as positive reinforcement. All procedures followed Canadian Council on Animal Care guidelines and were carried out at either McGill University or Western University and were approved by the respective University Animal Care Committees.

Hippocampal recordings were carried out using 1-4 high impedance (400-1500 kOhms) tungsten electrodes lowered each day to the right hippocampus, using co-registered image guidance for trajectory and depth, with examples seen in Figure 1e of recording locations. Most recordings were done in the mid to posterior putative CA3 region. Further information on electrode placement and targeting is available in Gulli et al. (2020). LPFC recordings were acquired using two 96-channel Utah arrays positioned at the posterior end of the principal sulcus, on the dorsal and ventral gyri of the principal sulcus and the anterior gyrus of the arcuate sulcus, spanning parts of area 8a and of area 9/46. The shank length was 1.5mm, and so was likely in layer II/III, and electrodes had an impedance ranging from 20 to 1500 kOhms. Signals were acquired at 30 kHz using one (HPC) or two (LPFC) 128-channel Cerebus recording systems (Blackrock Microsystems) and saved for later offline sorting, done with Plexon Offline sorter (version 4.5.0, Plexon Inc.). Time was not used as a feature during sorting, but units with waveform shapes that varied extensively or merged with other units were excluded from these analyses.

The tasks were very similar, taking place in a double ended Y maze, termed the Xmaze and described previously (Doucet, Gulli, & Martinez-Trujillo, 2016; Gulli et al., 2020). At the end of each Y, were two colored discs towards one of which the animals would navigate to receive the associated reward. The reward was dictated by the context, which was indicated by a texture that was applied to the walls, either a dark grey “steel” texture or a brown “wood” one. The highest value color in one context was the lowest in the other context (Fig.1d). The LPFC recordings were done with only this high and low option, but the HPC had a middle color that was worth half the reward in both contexts. Monkeys used the joystick to navigate to their chosen colour, receive the associated reward, and then turn around and navigate back towards the other end to make another choice. Figure 1 shows an example trial trajectory, and the trajectories for two example sessions (Fig. 1b,c).

### Behavioural Analysis

Monkeys were trained to be able to learn the task before recordings, and then presented with new combinations of colours each day, picked pseudo randomly to avoid a color occurring two days in a row. We used a performance analysis window of the 50 trials preceding the final 10 trials (excluded because performance may falter as satiation is reached).

### Calculating spike width

To separate neurons into putative principal cells and interneurons, we started by interpolating the waveform signals to 1MHz, and then aligning the waveforms for a neuron to the minimum of the trough. We then calculated the mean waveform and measured the duration between the minimum (trough) and the maximum (peak) in microseconds. To determine where to divide the neurons into narrow and broad spiking, for each area we fit two Gaussians, and used the local minimum as the threshold to separate them (HPC = 334µs, LPFC = 333µs). Because the results were within 1μs, we used the threshold for the LPFC as there were more neurons and it might be slightly more accurate. We then discarded all narrow neurons (<10%) because we did not have enough to analyse separately and focused our results on the putative pyramidal cells for the rest of the study.

### Calculating burst propensity

The initial analysis of ISIs was just done by taking all spikes recorded from broad spiking neurons during the task and pooling them for each area. We removed all ISIs greater than 60ms and then normalized histograms to value at the stable period at 60ms for each area. To assess the stability of the curve of the population, we calculated the difference between each point and plotted it for the HPC. We calculated the burst fraction as the fraction of all ISIs during the task that were equal to or below 7ms. Because this could start to correlate with high firing rates, we also made a burst index (BI) for the neurons based on the ISI histogram and the predicted ISI distribution based on a Poisson distribution with the calculated firing rate. To predict the probability of ISIs of a certain duration, we followed the method of Livingstone et al. (Livingstone et al., 1996), taking the firing rate averaged over the whole task. Using equation 1, where λ = firing rate, and t = time bin. We calculated the probability for each 1ms time bin from 2-40ms, and then normalized these

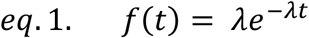

measures by the sum of all these predictions. We then did the same thing with the measured ISIs, normalizing by the sum of the measured ISIs between 2 and 40ms. We then summed the predicted values from 2 to 7ms and subtracted that from the sum of the measured values between 2 and 7ms. We divided this difference by the sum of the two sums to bind our index between −1 and 1 (see eq. 2). We then repeated this for threshold values of 4, 10, 15 and 20ms.

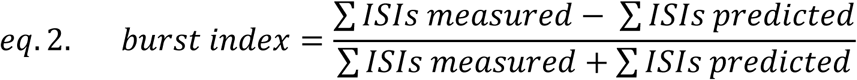

To calculate the *d’* noise measurement, we measured the mean and the variance at three locations, the trough (minimum peak), the peak (maximum peak), and the first point at the baseline of the spike, before any spike deflections, to get a measure of the noise. We then calculated two *d’* values, one comparing the trough to the baseline, and the other comparing the peak to the baseline both using equation 3. We then summed these two values and calculated the correlation with the BI (calculated at 7ms threshold).

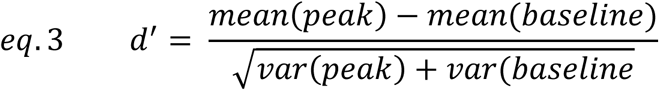

To calculate the geometric coefficient of variation (GCV) we first analysed only ISIs below 40ms, using equation 4. We used the GCV instead of the CV because the distribution of ISIs was not normal and was more similar to log-normal.

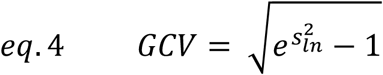

### Performance slope analysis

To assess performance, we chose 5 epochs: the first 20 trials, the last 20 trials, and the 20 trials centered on the ¼, ½, and ¾ marks of the session. We simply calculated the hit rate for the sessions during these epochs, and then performed a regression to determine if there was a positive slope, indicating that hit rate increased over the course of the session. We included 15 other sessions (Monkey T = 8, monkey B = 7) for the performance to show that the performance was consistent but did not analyse any neural data from these sessions.

### Burst and firing rate slopes

Similar to the performance analysis but restricted to periods after correct trials to control for reward effects. To calculate the rates of bursts or spikes, and to be able to use only correct trials that approximated the trials used to assess performance, we only used 10 correct trials: the first ten, the last ten, and the ten centred on the ¼, ½, and ¾ marks of the session. We only analysed broad spiking neurons that had at least one burst in at least two epochs, analysing the same neurons for both the burst and spike rates. To normalize the values, we summed the rates across epochs, and used this value to divide each epoch’s rate. We again ran a regression to determine if there was a consistent trend in the rates of bursts or spikes.

### Calculating information

To calculate mutual information, we used only correct trials. Bursts were detected as any group of three or more spikes where all ISIs were equal or less than 7ms. The timing of the burst was the onset of the first spike, and which ever period the first spike occurred in was considered the period that the burst happened in. We split the task into four behaviourally separate periods: the post reward period, the context period, the decision period, and goal approach period. The post reward period started at the end of the reward administration and is potentially when the monkey is incorporating the knowledge gained from the previous trial. This continues until they navigate to the start of the corridor where the context appears, which is the start of the context period. The decision period starts at the appearance of the target objects, when they must choose between the two, and the goal approach period starts at the beginning of the first turn of more than 10 degrees towards an object, and proceeds until they reach the target, just before reward administration. The period of reward administration was not analysed. The durations of each task period vary based on which period it is, and by trial, but we calculated rates based on the duration of each individual period. We only analysed broad spiking neurons, and separated neurons into high bursting neurons (BI>0, HBNs) and low-bursting neurons (BI<0, LBNs) and analysed these subpopulations separately. We used 60 sample rates from each trial period for each neuron to calculate the mutual information using the Neuroscience Information Theory Toolbox (Timme & Lapish, 2018). We then ran 5000 shuffles to generate the *p*-value for the mutual information. We then ran this whole process, starting from the subsampling, 50 times and analysed the means of the mutual information and the *p*-value for each neuron.

We also repeated this whole analysis with burst thresholds set at 10ms,15ms and 20ms, and fit a line to the 4 proportions of units that had significant burst rate information to determine whether this proportion increased with the threshold for the different subpopulations.

### Decoding analysis

For the decoding analysis, we used support vector machine (SVM) decoding from LIBSVM v.3.23 using five-fold cross-validation and a linear kernel on rates that were normalized between 0 and 1. We only used broad spiking, HBNs and again randomly sampled 60 rate pairs from each trial period and ran the decoding analysis on each set of rates before shuffling the labels and decoding again to get a measure of chance decoding. We repeated this 50 times to take the mean accuracy and mean chance decoding of the same trials for both spike rate and burst rate decoding. Significance was determined by permutation test, where the mean had to be greater than 95% of the shuffled performances to be significant. To examine the differences between the performance of the burst and spike decoder within an area, and to compare this across areas, we created a decoding index (see equation 5). This gave us 50 index values to run a t-test on.

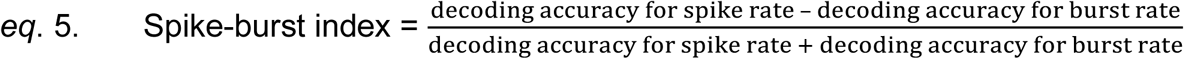

To analyze navigation and task parameter signals, we used the decision period of correct trials, and separated them into left and right decisions, and based on the two colours chosen. This gave us 4 combinations, and we used a subsample of 30 trials from 30 neurons, and we again used 5-fold cross validation. We ran this process 50 times on different subsamples and reported the distributions of mean decoding accuracies achieved for both areas for bursts and spikes.

### Optimized Decoders

For the optimized ensembles, we used the same set up for the SVM, but for the pool of neurons we only used HBNs that had significant mutual information (*p* < .05) for a specific rate. This limited the number of neurons we could use. To control for differences in number of eligible neurons in the two areas, for an iteration of the optimized decoder, we would randomly select up to 30 neurons from which we would build the decoder. For the first test, we used only neurons with significant mutual information for burst rate, which limited us to only 15 cells in LPFC, so we used this population size for both areas. Each neuron was tested individually to find the neuron with the best decoding accuracy. This neuron was then paired with every other neuron, and we ran an SVM on each duo to find the best duo. The best duo was used to find the best trio that included the best duo, and so on. This does not necessarily find the absolute optimal duo or trio, but it is an effective method for exploring the decoding space without having to exhaustively try all possible permutations, which can be computationally expensive. Indeed, it results in better decoding accuracies than simply using the best neurons based on individual performance, as illustrated in Leavitt et al (2017). After building the decoder to either 12 units for the initial analysis, or 20 units for the time window analysis (described below) we then selected another set of random units and repeated the process to build another optimized decoder, building 50 optimized decoders in total to give us a population of results to analyse.

For the decoders built to assess the contribution of neurons we only used neurons with significant information in the burst rate. We fit an exponential function (eq. 6) to each performance curve, and calculated the point where performance was 95% of the asymptote which is defined as 1-c. This gave us 50 points for each set of decoders so we could run t-tests on these points within areas and within spike and burst rates, however, we discarded any ensembles that did not reach the 95% point within 3 standard deviations of the distribution of 95% points.

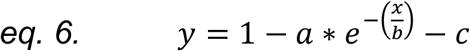

The time window analysis used different time windows within which to calculate the rates. Because we wanted to analyze the compression of the full signal, we only used the firing rates for this analysis, and used neurons that had significant firing rate mutual information values. We used time windows of 25, 50, 100, 150, 200, 300, 400, and 500 milliseconds instead of integrating over the whole trial period. We chose to center the windows in the middle of the period. We built the optimized decoders with pools of 30 neurons, but only built them to 20 neurons because they would have already saturated decoding performance before then. We built 50 optimized decoders for each time window for each area. To fairly compare the effect of time windows, we took the maximum of the average performance across ensemble sizes of the best decoder (500ms) and calculated the performance of all the decoders, at each ensemble size, as a fraction of such maximum performance. To compare these performance fractions, we took the HPC fractions for each time window and sample size and subtracted the corresponding LPFC fractions, which gave us a population of differences at each ensemble size and time window. We ran a two-way ANOVA on these differences to assess for effects of time window size or ensemble size on the differences in performance. To determine which differences between the two areas were significantly different from 0, we calculated a one-sample *t*-test for each ensemble size and time window.

### Working Memory task analysis

The recording and behavioural set-up for this task was the same, however the virtual environment navigated was different. It consisted of a circular arena with a 3×3 array of potential target locations, and a starting area on the side from where the monkey started each trial. Trials started with a Cue period where one of the 9 locations had a red fog presented for 3 seconds, followed by a 2 second Delay period, after which there was a 10 second Response period where the monkey had to navigate to the cued location. More information on task performance is included in (Roussy et al., 2021). We analysed 2 sessions from monkey T and one form Monkey B and combined the neurons from each to analyse burst indices and decoding performance. We were unable to record any hippocampal data in this task. We calculated the burst index as indicated above, and then ran a Kolmogorov-Smirnov test to determine whether there was a difference between the BI for the learning task and the working memory task. We also ran a decoder on bursting and firing rates during the Cue and Delay periods to determine if there was encoding of the target column location (grouped into three columns, left, right and centre).

## Supplementary figure captions

**Supplementary figure 1. Most neurons recorded in both areas were broad spiking putative principal cells**. a) distribution of HPC spike widths measured from trough to peak. The distribution is well fit by the sum of two gaussians, and we used the local minimum as the threshold to separate narrow and broad neurons. b) same as ‘a’ but for LPFC.

**Supplementary figure 2. Burst index values for different thresholds**. a) burst threshold distributions for HPC neurons. While the median values are similar for thresholds above 4ms, the 4ms threshold has very few neurons with positive values, and has a median that is far below the other medians. b) LPFC distributions of BI values. Again, the 4ms threshold median is much lower than the other medians which are otherwise tightly distributed.

**Supplementary figure 3. Burst performance in HPC is not a result of low signal-to-noise, but might be inflating LPFC performance**. a) Diagram explaining how d’ was calculated, comparing the difference between baseline before the spike to the trough and peak (see methods). b) BI values are not correlated with d’ in HPC. c) Slight negative correlation for LPFC neurons between BI and d’, suggesting that some bursts might be the result of noise entering the signal, however the high BI neurons do not have the lowest d’ values, which are concentrated below BI=0.

**Supplementary figure 4. HPC bursts correlate with performance**. a) Normalized burst rates during correct trials correlated with the hit rate for the HPC. b) Normalized spike rates did not significantly correlate with the hit rate. c&d) Same as a&b, except neither burst nor spikes have a significant correlation.

**Supplementary figure 5. Mutual information figures including the 4ms threshold**. a) Proportions of neurons with significant burst information using different thresholds. 4ms appears to be an outlier for the high bursting neurons, but not the low bursting ones. b) Mutual information values for neurons with significant information at different burst thresholds, as well as spikes.

**Supplementary figure 6. HPC and LPFC have similar distributions of silent periods**. Population density histograms for proportion of periods without any of the measured activity, bursts or spikes for HPC and LPFC. There was no significant difference between the distributions for bursts, but HPC has significantly higher proportions of silent periods with no spikes than the LPFC.

**Supplementary figure 7. HPC burst and spike codes and LPFC spike codes can decode both colour and side chosen during the goal approach period**. a) Decoding accuracy for burst and spike codes for which colour and which direction was chosen during the goal approach period. HPC burst and LPFC burst accuracy is not significantly different from chance. b) Average confusion matrices for the decoders in ‘a’. These allow visualisation of biases, particularly for side for the spiking decoders, where right colour 1 is often confused with right colour 2, but not with either combination of colour and left. The same is true for colours on the left which creates 4 larger squares of 4. This suggests that side is a stronger signal in the spike rate than colour, but colour is still being decoded, as indicated by the highest values in each column almost always being on the diagonal.

**Supplementary Figure 8. Foveation durations were similar between monkey pairs, suggesting that there were similar saccade rates**. a) The duration of foveations, or intersaccadic intervals for the NHPs used in either the HPC or LPFC recordings show a very similar distribution. b) The Empirical Cumulative Density Function (EDCF) of the foveation durations show there is very high overlap, with a very low Kolmogorov-Smirnov D of .05, which does reach significance due to the high power of this analysis with many thousands of foveations across many sessions.

