## Supplemental figures for "Distinct neural codes in primate Hippocampus and Lateral Prefrontal Cortex during associative learning in virtual environments"

### Supplementary Figure 1

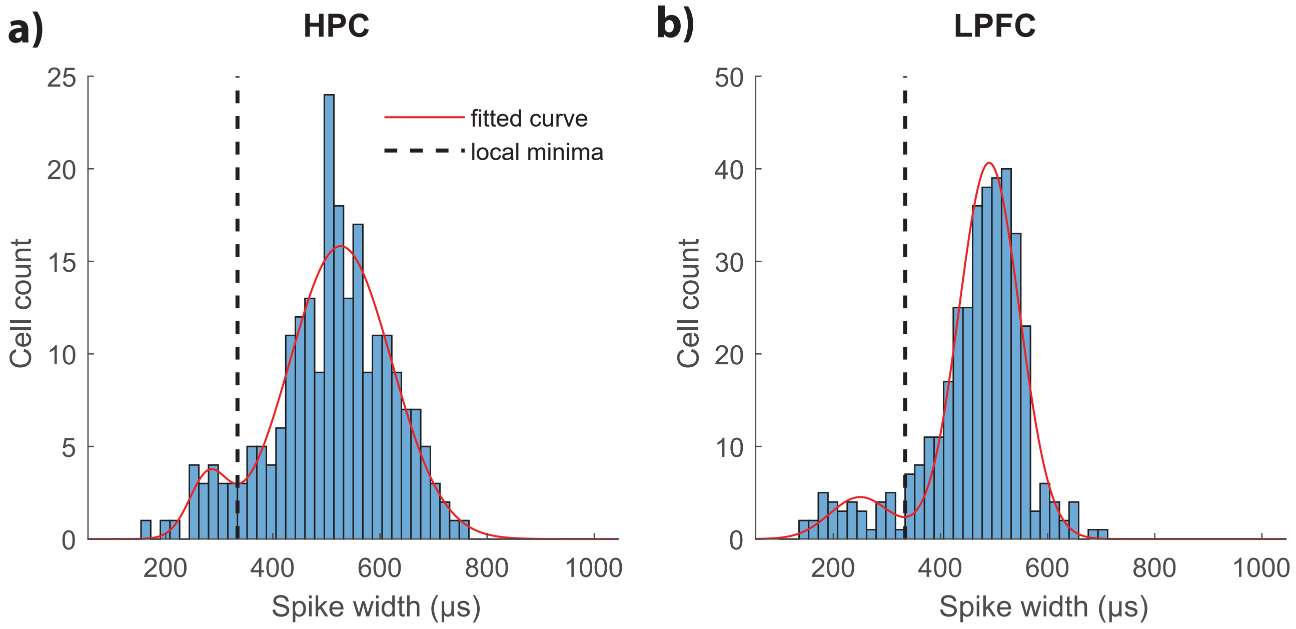

### Supplementary Figure 2

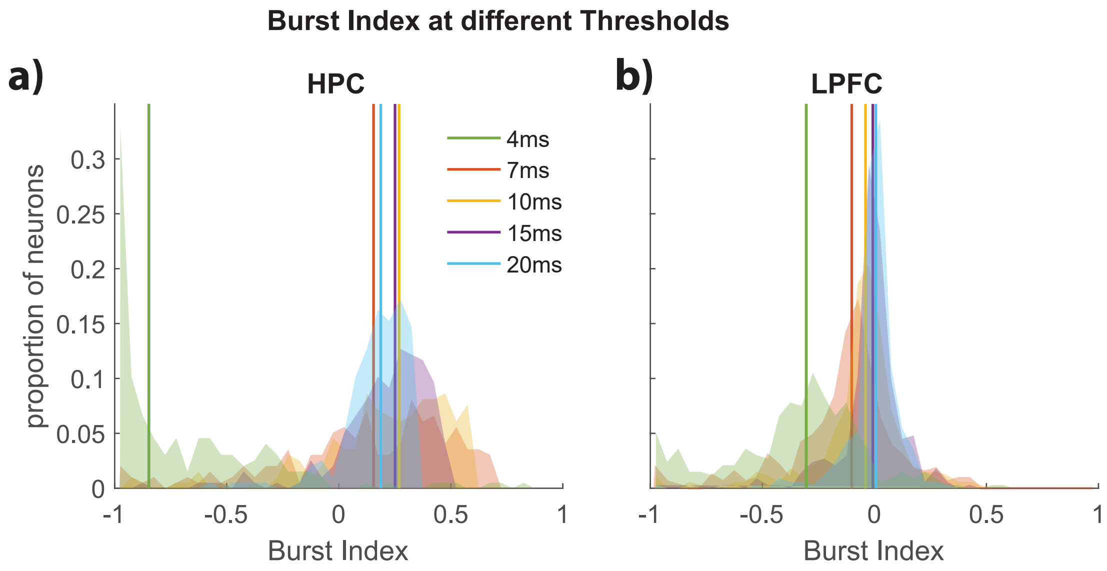

### Supplementary Figure 3

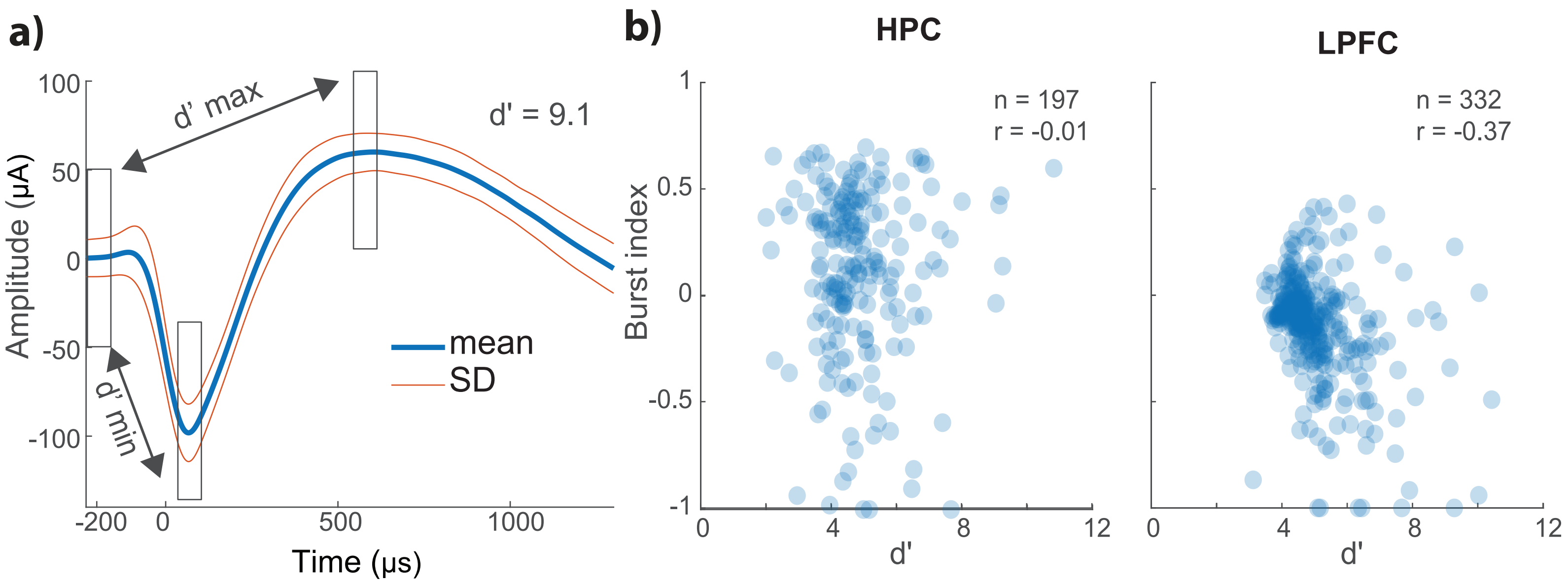

### Supplementary Figure 4

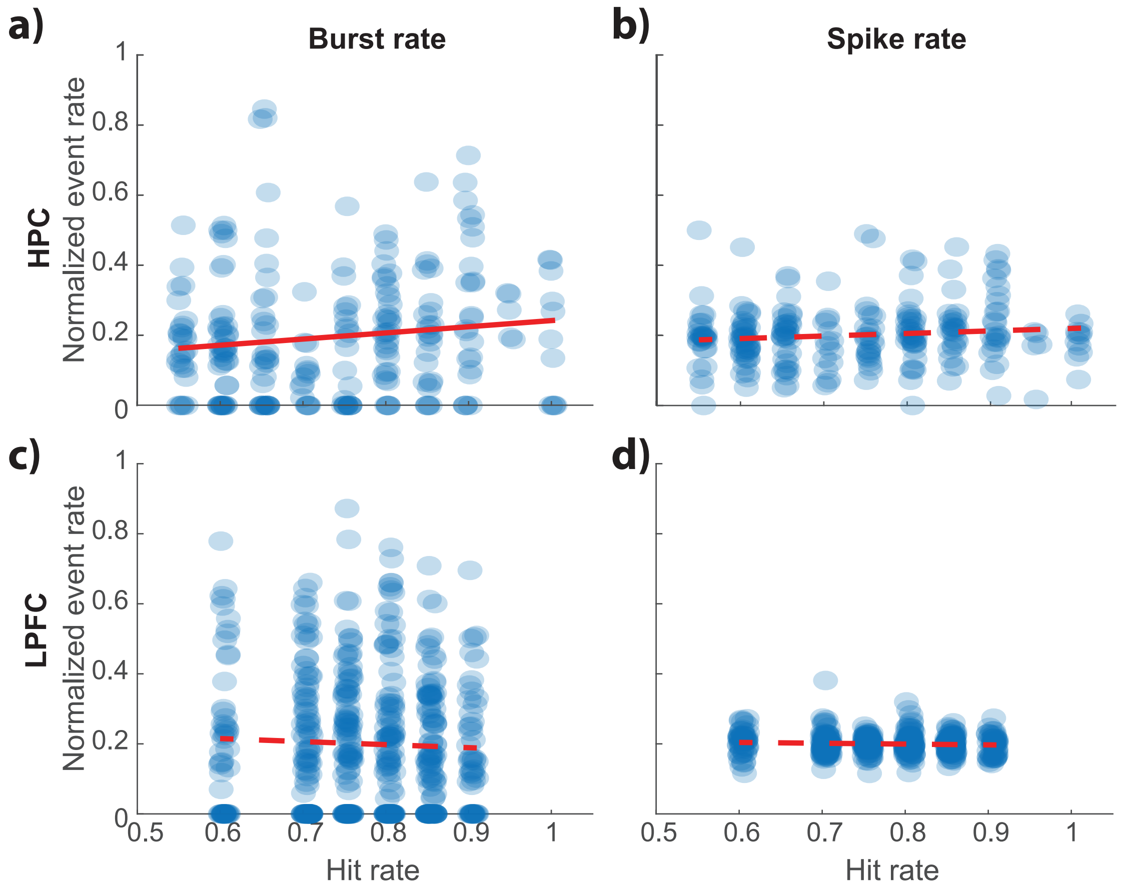

### Supplementary Figure 5

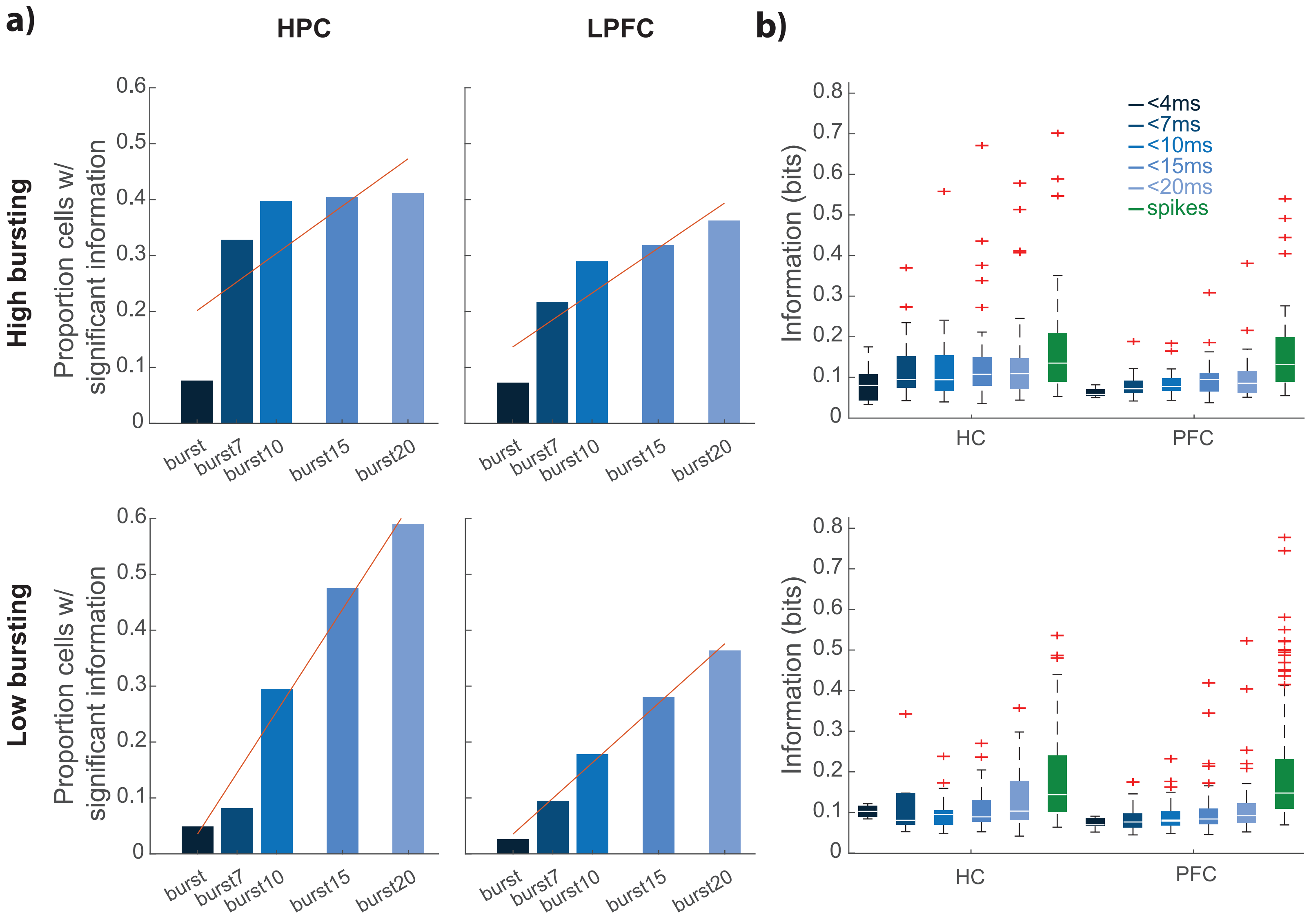

### Supplementary Figure 6

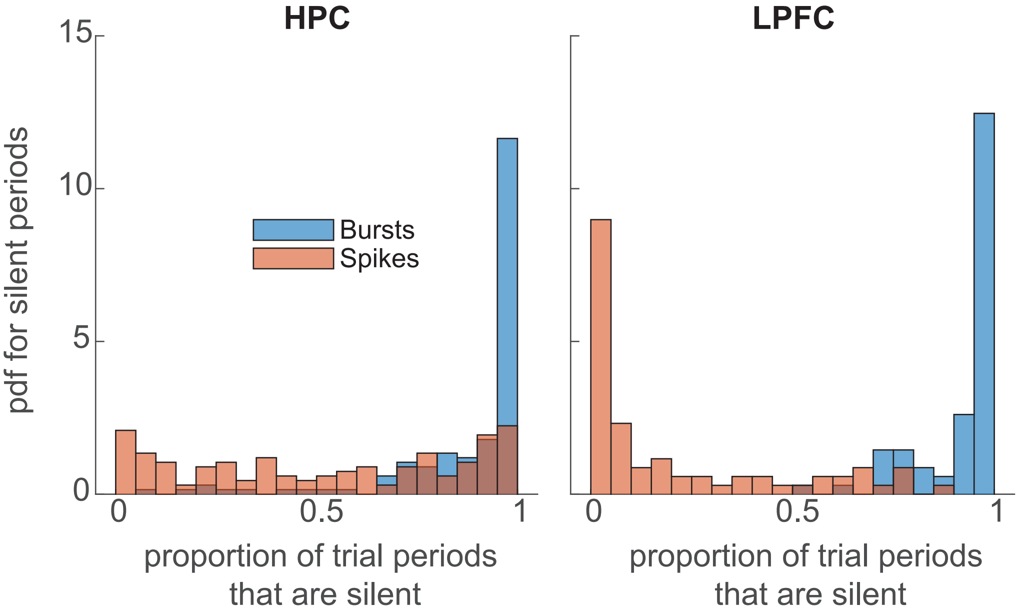

### Supplementary Figure 7

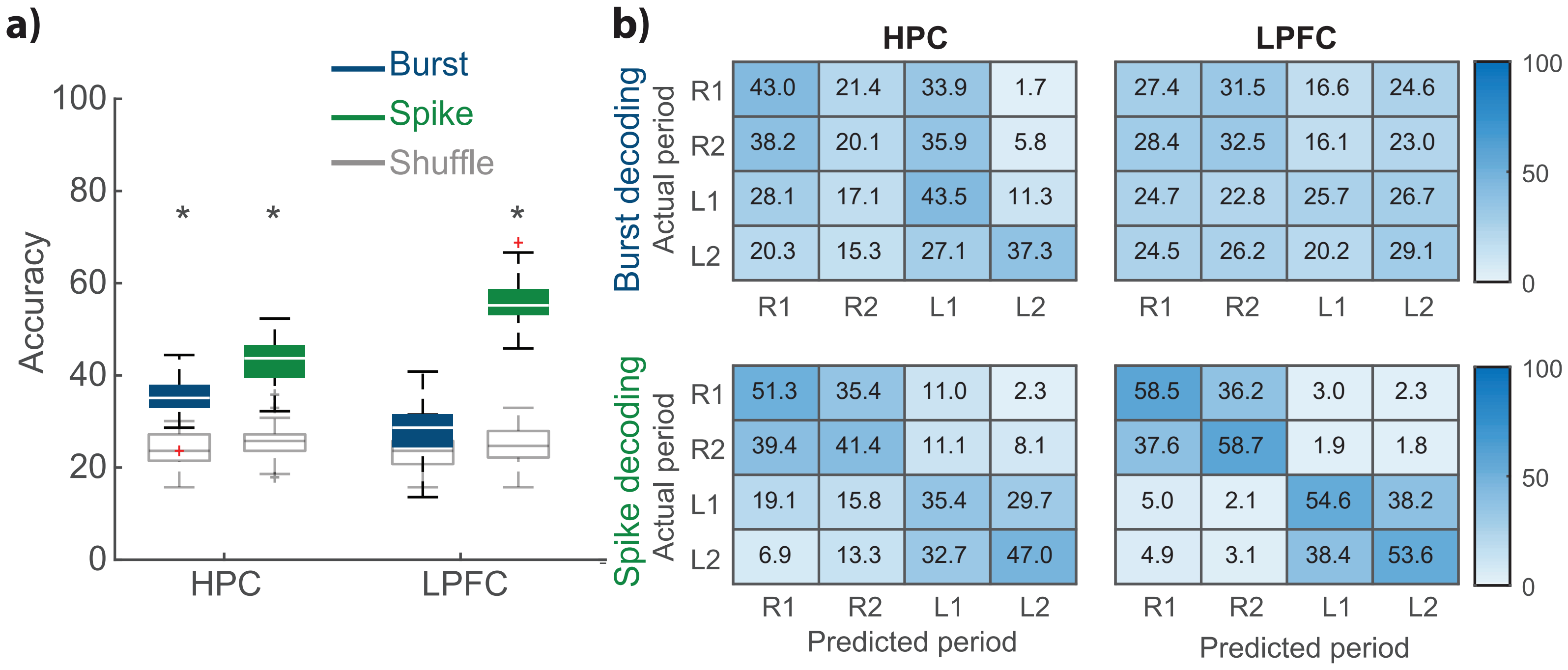

### Supplementary Figure 8

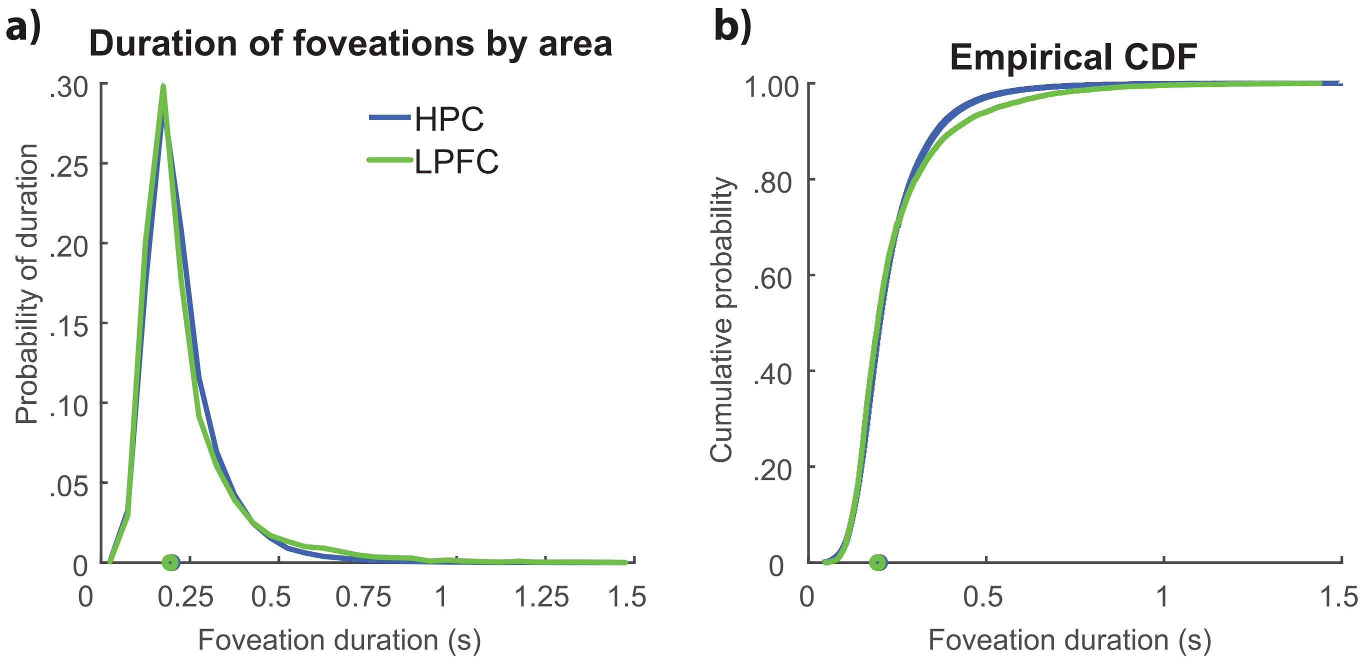
